# A Novel Zebrafish Luminescent Biosensor for Kidney Tubulopathy, Metal Toxicity, and Drug Screening

**DOI:** 10.1101/2025.09.19.677027

**Authors:** Han Lai, Monica Goldade, Svenja Keller, Isabelle Worms, Alessandro Luciani, Stephan C.F. Neuhauss, Vera I. Slaveykova, Olivier Devuyst, Zhiyong Chen

**Affiliations:** Department of Physiology, University of Zurich, Zurich, Switzerland; Department of Nephrology, First Affiliated Hospital of Chongqing Medical University, Chongqing, China; Department F.-A.Forel for Environmental and Aquatic Sciences, School of Earth and Environmental Sciences, Faculty of Science, University of Geneva, Geneva, Switzerland; Department of Molecular Life Sciences, University of Zurich, Zurich, Switzerland

**Keywords:** Proximal tubule, low-molecular-weight proteinuria, NanoLuc luciferase, receptor-mediated endocytosis, lysosome

## Abstract

An efficient endo-lysosomal pathway is crucial to mediate the reabsorption and processing of ultrafiltered solutes including low-molecular-weight (LMW) proteins by the epithelial cells lining the proximal tubule (PT) of the kidney. The zebrafish pronephros is increasingly used as a model system for congenital or acquired disorders that impair the endocytic uptake in PT cells, that cause an inappropriate loss of solutes and LMW proteins in the urine. Here, we describe a new reporter zebrafish line, termed *½vdbp-NanoLuc*, in which the vitamin D-binding protein is coupled to NanoLuc luciferase, for rapid, large-scale detection of PT dysfunction and LMW proteinuria. We demonstrate the reliability and value of the *½vdbp-NanoLuc* biosensor in fish models of monogenic endolysosomal diseases, gentamicin and cisplatin-induced nephrotoxicity, and metal contamination. This novel system provides mechanistic insights into the cadmium- and copper-induced PT dysfunction and, when combined with a swimming test, a platform for drug screening to alleviate cisplatin toxicity.

## INTRODUCTION

The epithelial cells lining the proximal tubule (PT) of the kidney play a key role in controlling body homeostasis through their capacity to reabsorb a large variety of solutes. The latter include low-molecular-weight (LMW) proteins such as hormones, vitamins and their binding proteins, enzymes, immunoglobulin light chains, as well as drugs and toxins which are ultrafiltered but not/minimally excreted in the final urine. Receptor-mediated endocytosis and autophagy are critical to mediate the reabsorption and processing capacity of PT cells (van der Wijst et al., 2019). Congenital or acquired disorders, as well as exposure to drugs and toxins may impair the endolysosomal machinery and the reabsorption capacity of PT cells, causing epithelial cell dysfunction and inappropriate loss of LMW proteins and vital nutrients into the urine (Devuyst and Luciani, 2015; van der Wijst et al., 2019). The kidney PT is also the main site of accumulation and toxicity of metals including cadmium, copper, mercury, and lead, which are extensively used in agriculture and industrial activities and significantly contribute to environmental pollution (Lentini et al., 2017; Rawee et al., 2023). Clinical and experimental studies have demonstrated that the detection of LMW proteinuria is the most consistent and sensitive approach to detect PT dysfunction (Bernard and Lauwerys, 1995; Bokenkamp, 2020; Devuyst O, 2018; Hall et al., 2022).

The zebrafish has emerged as a prominent model for studying human kidney disease and chemical nephrotoxicity due to its highly conserved nephron segment patterning (Outtandy et al., 2018; Poureetezadi and Wingert, 2016). In particular, the PT cells of the zebrafish pronephros express the endocytic receptors megalin and cubilin, mediating a high endocytic uptake - similar to that observed in the human kidney. The generation of zebrafish models of endolysosomal disorders including Donnai-Barrow syndrome (Veth et al., 2011), Lowe syndrome (Oltrabella et al., 2015) and cystinosis (Festa et al., 2018) validated the use of this model system to investigate congenital disorders of the kidney proximal tubule.

Standard assessment of PT function in zebrafish previously relied on visualizing the uptake of injected LMW fluorescent tracers. These assays are labor intensive, challenging to quantify, and limited by lack of sensitivity, elevated background levels, and difficulty to perform accurate and reproducible intravenous tracer injection. These limitations render injection-based assays not appropriate for large-scale assessment of PT dysfunction in the context of drug discovery and toxicology screens. To overcome this barrier, we previously developed a transgenic *½vdbp-mCherry* reporter zebrafish line for monitoring LMW proteinuria based on the quantification of the vitamin D-binding protein (VDBP)-mCherry in the urine by ELISA (Chen et al., 2020). This model system was validated as a consistent tool to assess PT dysfunction in endolysosomal disorders (Chen et al., 2020). Yet, the scope and applicability of this ELISA-based detection system is limited due to the narrow dynamic range, cost and time-intensive nature of the assay.

Luminometry is increasingly used to overcome the limitations of ELISAs, due to its higher sensitivity, broad dynamic range, and simpler handing with no washing steps (Hall et al., 2021; Jackson et al., 1996). Here, we applied the bioluminescence concept to establish a new *½vdbp-NanoLuc* zebrafish biosensor for LMW proteinuria, by coupling VDBP to NanoLuc luciferase, the catalytic subunit of the luciferase isolated from *Oplophorus gracilirostris* (Inouye et al., 2000). We validated the bioluminescent biosensor using genetic models of endolysosomal diseases (Chen et al., 2020; Festa et al., 2018). and nephrotoxins known to induce PT dysfunction, including metals (Hentschel et al., 2005; Kim et al., 2020; Rawee et al., 2023). We showed the value of this new line for drug screening to alleviate lysosomal defects and PT dysfunction associated with cisplatin toxicity.

## RESULTS

### Establishment of the ½vdbp-NanoLuc zebrafish reporter system

The novel bioluminescent reporter line is based on VDBP, a relevant marker of LMW proteinuria, coupled to NanoLuc luciferase. The expression vector included the 24 kDa, 207 N-terminal amino acids of zebrafish vdbp tagged with 19 kDa NanoLuc luciferase (termed vdbp-NanoLuc) for expression in the liver under control of the liver-type fatty acid binding protein *lfabp10a* promoter and the transgenesis marker *cmlc2::EGFP* to express EGFP in the heart (Fig. 1A), allowing to sort the early embryos by EGFP fluorescence (Huang et al., 2003). The resulting transgenic *Tg(lfabp::½vdbp-NanoLuc/cmlc2::EGFP)* zebrafish larvae (termed *½vdbp-NanoLuc* line) expresses the 43 kDa vdbp-NanoLuc in the liver (Fig. 1B) and EGFP in the heart (Fig. 1C).

**Fig. 1.**
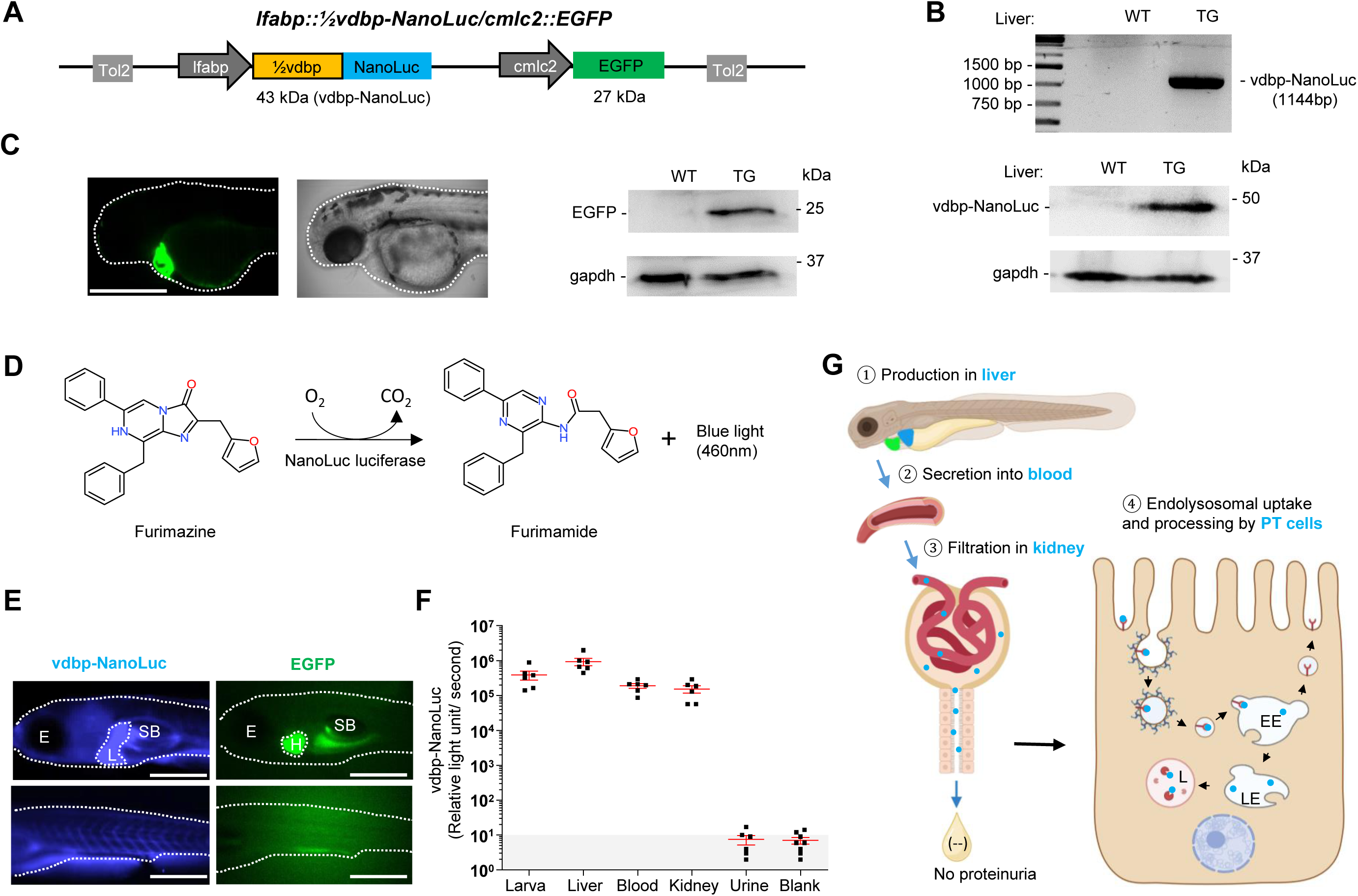
Establishment of the ½vdbp-NanoLuc zebrafish reporter system. (A) Construction of plasmid vector *lfabp::½vdbp-NanoLuc* containing the transgenesis marker *cmlc2*:*:EGFP* and Tol2 transposon elements. The N-terminal 207 a.a. of zebrafish vdbp (½vdbp) are fused with NanoLuc luciferase, which is predicted to have a molecular weight of 43 kDa. *lfabp* promoter drives the expression of vdbp-NanoLuc in the liver and *cmlc2* promoter controls the expression of EGFP in the heart. Tol2: transposable elements facilitating the integration of plasmid DNA into the embryo genome. (B) Analysis of *½vdbp-NanoLuc* cDNA (1144bp) after amplification by Polymerase chain reaction and immunoblotting analysis of vdbp-NanoLuc in liver tissue. WT: wild type and TG: transgenic. (C) Observation of EGFP fluorescence in the heart of transgenic larvae visualized at 2 days post-fertilization (dpf) by Light sheet microcopy and detection of EGFP (27 kDa) by immunoblotting in larval lysate coming from wild type (WT) and transgenic *½vdbp-NanoLuc* (TG) larvae. Bar = 0.5 mm. (D) The oxidation of furimazine by NanoLuc luciferase in the presence of O_2_ produces furimamide, CO_2_ and blue light of wavelength 460nm. (E) Observation of *½vdbp-NanoLuc* larva at 5 dpf by bioluminescence microscopy using Olympus LV200. Incubating with its substrate furimazine, bioluminescence light from vdbp-NanoLuc (blue) was seen at least in liver and vessels, and EGFP fluorescence (green) was observed in heart. E: Eye, L: Liver, SB: swim bladder, H: heart. Bar = 0.5 mm. (F) Quantification of vdbp-NanoLuc luciferase activity in transgenic zebrafish by luminometry: whole larval lysates (lane 1), isolated liver (lane 2), blood (lane 3), isolated kidney (lane 4), urine collected from 5 dpf larvae (lane 5), blank (only E3 medium, lane 6); *n* = 6. (G) Schema illustrating the fate of recombinant protein vdbp-NanoLuc under normal condition: vdbp-NanoLuc is produced in liver ①, secreted into the bloodstream ②, passed through the glomerulus filtration in kidney ③, reabsorbed and processed by PT epithelial cells ④. EE: early endosome, LE: late endosome, L: lysosome.

The NanoLuc luciferase catalyzes the oxidation of furimazine and related substrates including Hikarazine Z108, yielding a blue light at 460 nm (Fig. 1D; Fig. S1A) (Coutant et al., 2020; England et al., 2016), with a light intensity directly proportional to the relative concentration of NanoLuc luciferase for both substrates (Fig. S1B,C). Under a bioluminescence microscope, blue light was observed in the liver and blood vessels of 5 dpf *½vdbp-NanoLuc* larvae in the presence of furimazine, while EGFP was in the heart (Fig. 1E). The vdbp-NanoLuc was detected in larval lysate as well as in liver, blood and kidney samples isolated from transgenic zebrafish (Fig. 1F). Urine collected from 5 dpf larvae did not show significant NanoLuc luciferase activity, confirming the complete reabsorption of vdbp-NanoLuc by PT cells under normal conditions (Fig. 1F,G). These data validate that the vdbp-NanoLuc protein produced in the liver of *½vdbp-NanoLuc* zebrafish is released in blood, filtered, and reabsorbed by kidney PT cells, so that it is absent from the urine much like natural VDBP under physiological conditions (Fig. 1G).

### Validation of the bioluminescent vdbp-NanoLuc reporter system in genetic models

We validated the vdbp-NanoLuc reporter system in two paradigmatic disorders affecting the receptor-mediated endocytosis and endolysosomal pathways, that consistently induce PT dysfunction and LMW proteinuria (Fig. 2A) (van der Wijst et al., 2019).We first analyzed *½vdbp-NanoLuc* zebrafish knock-out for *lrp2a* gene (*lrp2a KO*), which encodes the endocytic receptor megalin (Saito et al., 1994). The *lrp2a KO* zebrafish have enlarged eye globes in adult (Fig. 2B), mimicking the ocular anomalies in patients affected by Donnai-Barrow syndrome due to *LRP2* mutations (Kantarci et al., 2007). The defective endocytosis in *lrp2a*-deficient zebrafish is reflected by the highly reduced number of endocytic vesicles in the subapical compartment of PT cells analyzed by transmission electron microscopy (TEM) (Fig. 2C).

**Fig. 2.**
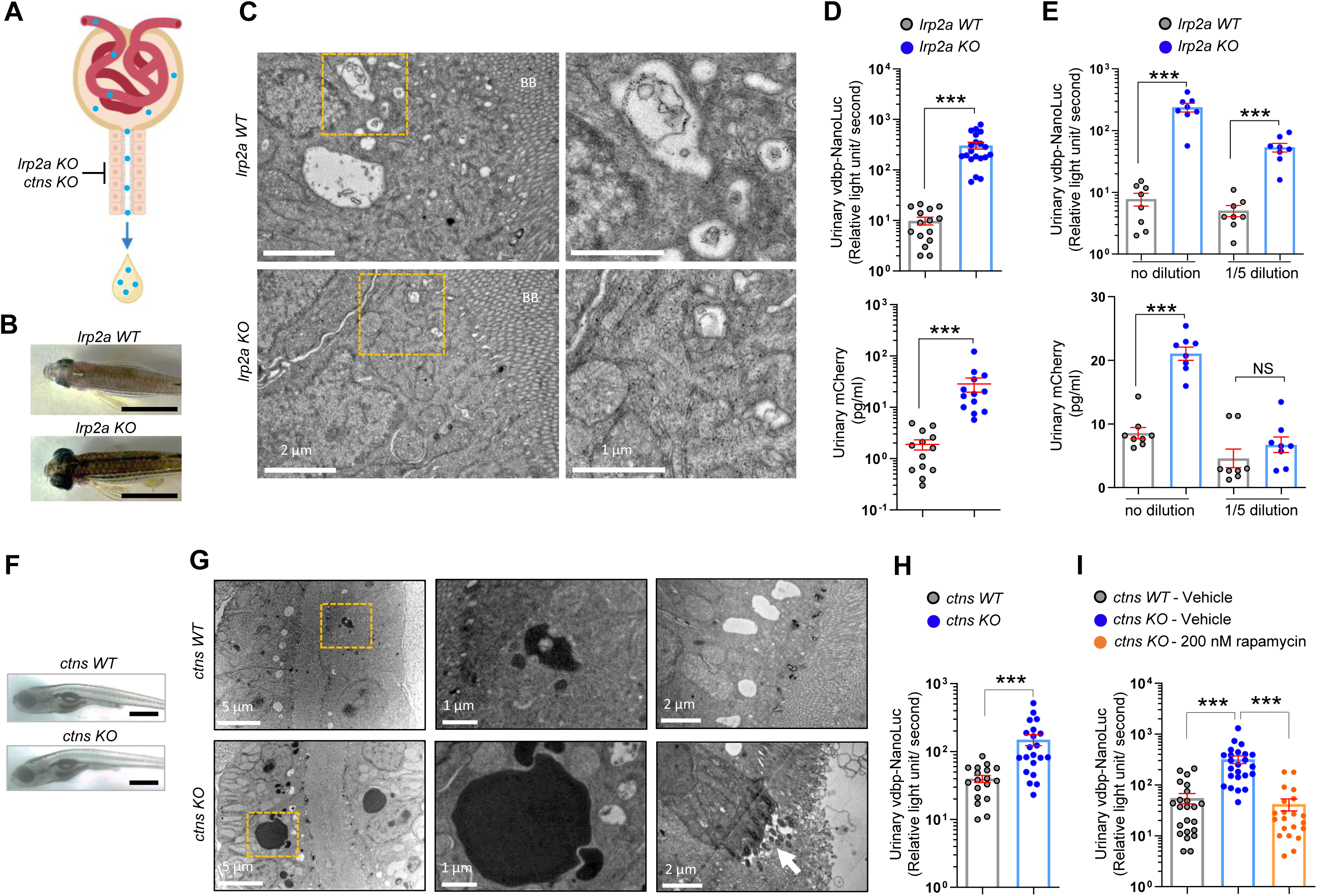
Validation of the bioluminescent vdbp-NanoLuc reporter system in genetic models. (A) Schema illustrating the genetic models of PT dysfunction caused by inactivation of megalin/lrp2a receptor (*lrp2a KO*) or deficiency of lysosomal cystine transporter cystinosin/ctns (*ctns KO*). (B) Dorsal views showing enlarged eye globes in *lrp2a KO* adult zebrafish. Bar = 1 cm (C) Transmission electron microscopy showed loss of endocytic vesicles in PT of *lrp2a KO* zebrafish larvae. Yellow squares contain images at a higher magnification; BB: brush border; Bar = 2 µm (left panel), Bar = 1 µm (right panel). (D) Overnight urine were collected from *lrp2a WT* and *lrp2a KO ½vdbp-NanoLuc* or *½vdbp-mCherry* zebrafish larvae at 5 days post-fertilization (dpf), and analyzed by luminometric assay or mCherry ELISA; *½vdbp-NanoLuc*: *n* = 15 (*lrp2a WT*) and *n* = 20 (*lrp2a KO*); *½vdbp-mCherry*: *n* = 13. (E) Transgenic *lrp2a* mutant *½vdbp-NanoLuc* zebrafish was crossed with *lrp2a mutant ½vdbp-mCherry* zebrafish to produce double transgenic *lrp2a KO* larvae expressing both VDBP tracers. Urine samples were collected and analyzed without dilution (no dilution) or after 5x dilution in E3 medium (1/5 dilution) for both VDBP reporter proteins in the same urine sample. After 5x dilution, the urinary NanoLuc luciferase activity in *lrp2a KO* larvae is still significantly higher than those in *lrp2a WT* larvae, while no difference was observed between the two groups when urine samples were analyzed by mCherry ELISA. *n* = 8. (F) Morphology of *ctns KO* larvae at 14 dpf. Bar = 1 mm (G) Representative micrographs showing the electron-dense vesicles and sporadic loss of brush border in the PT epithelial cells of *ctns KO* larvae at 14 dpf. Yellow squares contain images at a higher magnification; Arrow: loss of brush border; Bar = 5 µm (upper panel), Bar = 1 µm (middle panel) and Bar = 2 µm (bottom panel). (H) Quantification of vdbp-NanoLuc luciferase activity in urine samples obtained from 14-day-old *ctns WT* and *ctns KO ½vdbp-NanoLuc* zebrafish larvae. *n* = 18 (*ctns WT*) and *n* = 21 (*ctns KO*). The urinary vdbp-NanoLuc was slightly increased in transgenic *ctns WT* larvae at baseline once they are kept in fish facility water and under feeding. (I) *ctns KO ½vdbp-NanoLuc* larvae were treated with fish facility water containing vehicle or 200 nM rapamycin, followed by assessment of urinary vdbp-NanoLuc at 14 dpf. *n* = 23 (*ctns WT* -vehicle), *n* = 24 (*ctns KO* -vehicle) and *n* = 21 (*ctns KO* -rapamycin). Plotted data represent mean ± SEM. Nonparametric Mann-Whitney test, ***P < 0.001 relative to *lrp2a WT* or *ctns WT*; NS, non-significant.

Quantification of urinary vdbp-NanoLuc collected at 5 dpf revealed a major, 30-fold increase of bioluminescent signal in *lrp2a KO* compared to *lrp2a wild-type (WT)* larvae (306±56 vs. 9.8±1.8 RLU/sec, respectively; p < 0.001) (Fig. 2D upper panel). The increased VDBP signal in *lrp2a KO ½vdbp-NanoLuc* was twice more important than that observed when using ELISA in *lrp2a*-deficient *½vdbp-mCherry* larvae compared to *lrp2a WT* (28.4±8.7 vs. 1.9±0.4 pg/ml, respectively; p < 0.001) (Fig. 2D bottom panel), allowing a significantly higher sensitivity of detection upon dilution of the urine samples (Fig. 2E).

We next analyzed the bioluminescent VDBP signal in *ctns*-deficient *½vdbp-NanoLuc* zebrafish larvae (*ctns KO*). Inactivating mutations in *CTNS*, which encodes the lysosomal cystine transporter cystinosin, cause cystinosis - a lysosomal storage disorder causing PT dysfunction early in life. We previously demonstrated that *ctns KO* zebrafish larvae develop cystine accumulation, impairment of lysosomal degradation activity and autophagy, as well as LMW proteinuria (Berquez et al., 2023; Festa et al., 2018). No obvious morphological change was observed in *ctns KO*, as compared to *ctns* wild type (*ctns WT*) larvae (Fig. 2F). However, ultrastructural analysis by TEM confirmed the accumulation of large electron-dense vesicles and sporadic loss of the brush border in PT epithelial cells of *ctns KO* larvae (Fig. 2G). A major, 4-fold increase in the urinary vdbp-NanoLuc levels was observed in *ctns KO larvae* compared to *ctns WT ½vdbp-NanoLuc* larvae and (150±22 vs. 40.1±4.9 RLU/sec, respectively; p<0.001) (Fig. 2H). Cystine accumulation is known to induce mTORC1 hyperactivation in *ctns*-deficient larvae, which can be rescued with treatment of rapamycin, an inhibitor of mTORC1 (Berquez et al., 2023). Accordingly, we could observe a rescue of the abnormal urinary excretion of vdbp-NanoLuc with a treatment of 200 nM rapamycin in *ctns KO ½vdbp-NanoLuc* larvae, as compared to DMSO-treated *ctns KO* group (42.0±11.1 vs. 320±56 RLU/sec, respectively; p<0.001) (Fig. 2I).

These data show that the vdbp-NanoLuc reporter line can reliably and sensitively monitor LMW proteinuria in genetic models of PT dysfunction and is useful for testing therapies.

### Bioluminescent vdbp-NanoLuc reporter to detect drug nephrotoxicity

To determine whether the vdbp-NanoLuc reporter system can be used for monitoring drug-induced nephrotoxicity, we first exposed the *½vdbp-NanoLuc* larvae to gentamicin (Fig. 3A), an aminoglycoside antibiotic known to cause lysosomal phospholipidosis and PT dysfunction (Chen et al., 2020; Hentschel et al., 2005). In order to study the lysosome morphology and dynamics in zebrafish larvae, we employed the PT-specific reporter line *Tg(PiT1::ctns-EGFP)*, which expresses the lysosomal cystine transporter cystinosin/ctns labeled with EGFP in PT epithelial cells (Chen et al., 2020). Exposure to gentamicin induced the accumulation ctns-EGFP positive vesicles in *Tg(PiT1::ctns-EGFP) larvae*, with significantly enhanced fluorescent signal intensity (6.2±0.9 vs. 2.7±0.6, respectively; p < 0.01) and higher number of GFP-positive vesicles (102±11 vs. 46±7.3 respectively; p < 0.001) in PT cells, as compared to the vehicle-treated controls (Fig. 3B). Gentamicin exposure was also reflected by an accumulation of enlarged electron-dense lysosomal compartments in PT cells (Fig. 3C). Gentamicin induced a ∼100-fold increase in VDBP signal in *½vdbp-NanoLuc* zebrafish larvae compared to controls (756±216 vs. 7.9±1.0 RLU/sec, respectively; p < 0.001). The PT dysfunction caused by gentamicin toxicity was significantly rescued by co-treatment with taurine (gentamicin+taurine: 230±72 RLU/sec; p < 0.01 vs. gentamicin alone; Fig. 3D), as previously described (Chen et al., 2020).

**Fig. 3.**
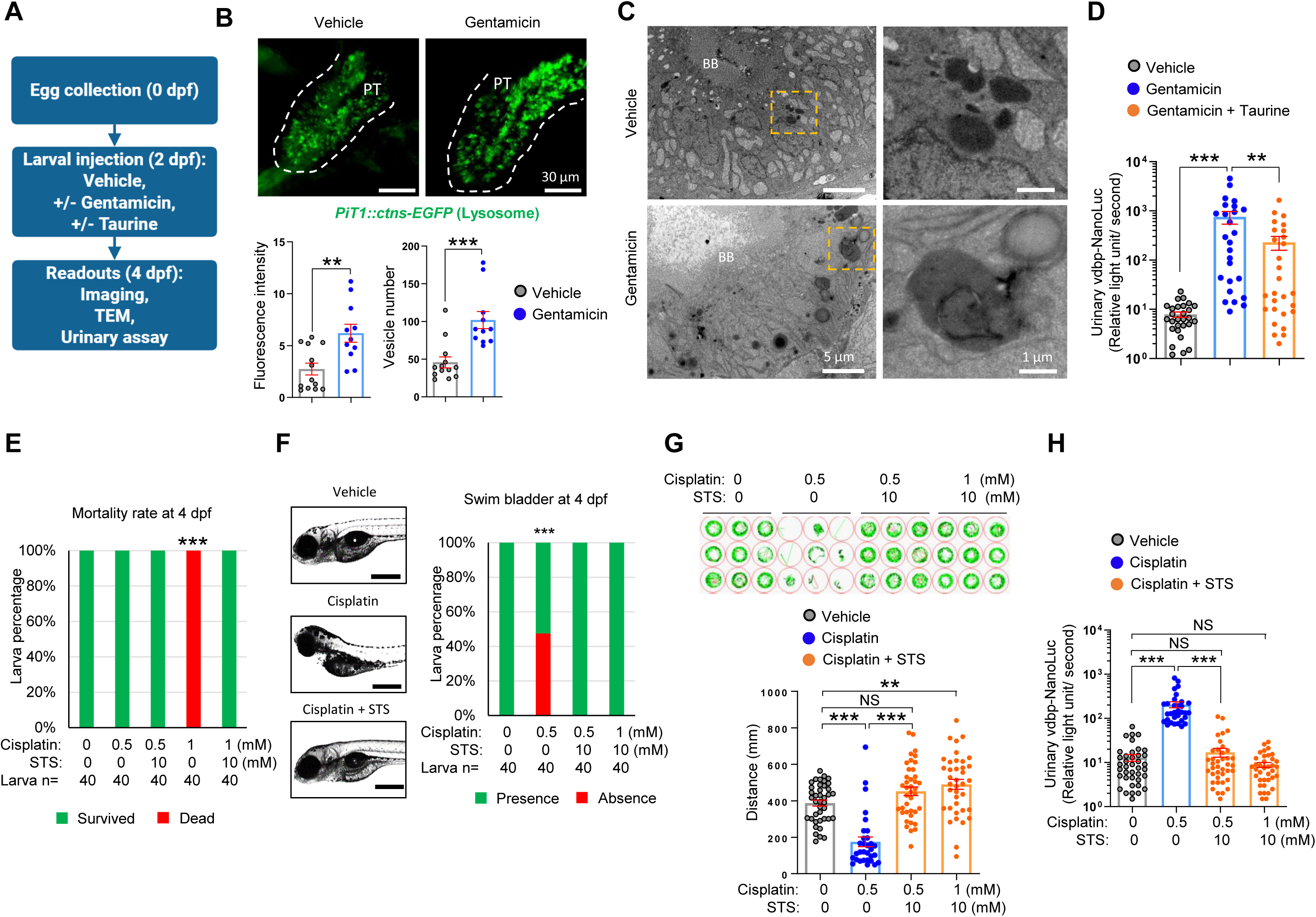
Bioluminescent vdbp-NanoLuc reporter to detect drug nephrotoxicity. (A) Experimental protocol of gentamicin nephrotoxicity assessment and its rescue by taurine co-treatment using transgenic zebrafish larvae. (B) Microscopic examination of 4-day-old gentamicin-treated *Tg(PiT1::ctns-EGFP)* zebrafish larvae expressing lysosomal marker ctns-EGFP in PT cells by multiphoton fluorescence microscopy. Accumulation of ctns-EGFP-positive vesicles were observed in PT of gentamicin-treated transgenic zebrafish larvae. PT: proximal tubule; *n* = 13 (vehicle), *n* = 11 (gentamicin); Bar = 30 µm. (C) Ultrastructure of pronephros in 4-day-old larvae treated by gentamicin or vehicle were analyzed by transmission electron microscopy. The electron-dense vesicles were accumulated in PT epithelial cells, which reflect the lysosomal phospholipidosis. Yellow squares contain images at a higher magnification; BB: brush border; Bar = 5 µm (left panel) and Bar = 1 µm (right panel). (D) Urine samples were obtained at 2 days post-injection from *½vdbp-NanoLuc* larvae treated with vehicle, gentamicin alone or gentamicin/taurine and analyzed by luminometry; The big variation in the gentamicin-injected group might be due to the fact that the drug is introduced into larvae by intravenous injection of nano-liter of solution, for which the injection volume is difficult to be accurate. *n* = 28 (vehicle), *n* = 26 (gentamicin) and *n* = 28 (gentamicin/taurine). (E) Larval lethality of 4-day-old larvae treated with vehicle, cisplatin (0.5 mM or 1 mM) alone or co-incubated with cisplatin (0.5 mM or 1 mM) /10 mM sodium thiosulfate (STS) for 24 hours. The treatment of 1mM cisplatin alone for 24 hours was lethal for zebrafish larvae, which was prevented by co-treatment with 10 mM STS; *n* = 40. (F) Morphological examination of 4-day-old larvae treated with vehicle, 0.5 mM cisplatin alone or co-treatment of cisplatin (0.5mM or 1 mM)/10 mM STS and semi-quantitative scoring of inflation defects of swim bladder; *n* = 40; Bar = 0.5 mm. (G) Behavioral analysis of 5-day-old zebrafish larvae incubated with vehicle, 0.5 mM cisplatin alone, or co-incubated with cisplatin (0.5mM or 1 mM)/STS after movement tracking with Zebrabox. *n* = 40 (vehicle), *n* = 31 (cisplatin alone), *n* = 40 (0.5 mM cisplatin + 10 mM STS), *n* = 38 (1mM cisplatin + 10 mM STS). (H) Overnight urine were collected from 5-day-old *½vdbp-NanoLuc* zebrafish larvae treated with vehicle, 0.5 mM cisplatin alone or co-incubated with cisplatin (0.5mM or 1 mM)/ 10 mM STS for 24 hours and analyzed by luminometry; *n* =39 (Vehicle), *n* = 31 (0.5 mM cisplatin alone), *n* = 39 (0.5 mM cisplatin + 10 mM STS), *n* = 37 (1 mM cisplatin + 10 mM STS).

Exposure of PT cells to the chemotherapeutic agent cisplatin results in cell injury and LMW proteinuria (Wilmes et al., 2015). In rodent models, cisplatin toxicity can be rescued by the addition of sodium thiosulfate (STS) (Dickey et al., 2005). Treatment of *½vdbp-NanoLuc* larvae with 1.5mM cisplatin for 6 hours impaired the inflation of swim bladder (Fig. S2A) and caused an abnormal urinary excretion of vdbp-NanoLuc (Fig. S2B), as described earlier (Chen et al., 2020). To optimize the cisplatin treatment protocol for drug screening, we tested lower concentrations of cisplatin for 24 hours. The 24-hour incubation of 2 dpf larvae with 0.5 mM cisplatin was non-lethal (Fig. 3E) but induced a 50% incidence of defective swim bladder inflation at 4 dpf (Fig. 3F). The latter resulted in impaired swimming capacity compared to vehicle treated larvae (175±26 mm vs. 387±17 mm, respectively; p < 0.001) (Fig. 3G). Administration of 10 mM STS rescued lethality (Fig. 3E) as well as swim bladder defects (Fig. 3F) and swimming performance (Fig. 3G) in cisplatin treated larvae. Exposure of *½vdbp-NanoLuc* larvae to 0.5 mM cisplatin induced a 16-fold increase in the VDBP signal in fish pool water, compared to vehicle-treated controls (205±32 vs. 12.9±2.2 RLU/sec, respectively; p < 0.001), with expected rescue by 10 mM STS co-treatment (17.0±3.8 RLU/sec; p < 0.001 vs. cisplatin alone) (Fig. 3H). Similarly, no VDBP signal was detected in larvae co-treated with 1 mM cisplatin and 10 mM STS (Fig. 3H). Collectively, these data demonstrate that the bioluminescent vdbp-NanoLuc reporter system can reliably monitor nephrotoxin-induced PT dysfunction and potential therapies that alleviate drug/chemical nephrotoxicity.

### vdbp-NanoLuc reporter as biosensor for detecting metal-induced nephrotoxicity

Environmental exposure to metal species resulting from agriculture and industrial activities represents a global health concern (Lentini et al., 2017; Rawee et al., 2023). The ability of the kidney PT to reabsorb and concentrate metals including cadmium/Cd, copper/Cu and lead/Pb makes it a primary target for metal-induced toxicity. To test the potential value of the vdbp-NanoLuc system in this context, we exposed zebrafish larvae from 2 dpf to Cd, Cu and Pb at environmentally relevant concentrations (from 1 to 100 µg/L, molar concentration is used in the following text), and analyzed for larval metal quantity, swim bladder inflation, swimming behaviors and urinary vdbp-NanoLuc at 5 dpf, kidney ultrastructure and expression of fluorescent reporter proteins (Fig. 4A).

**Fig. 4.**
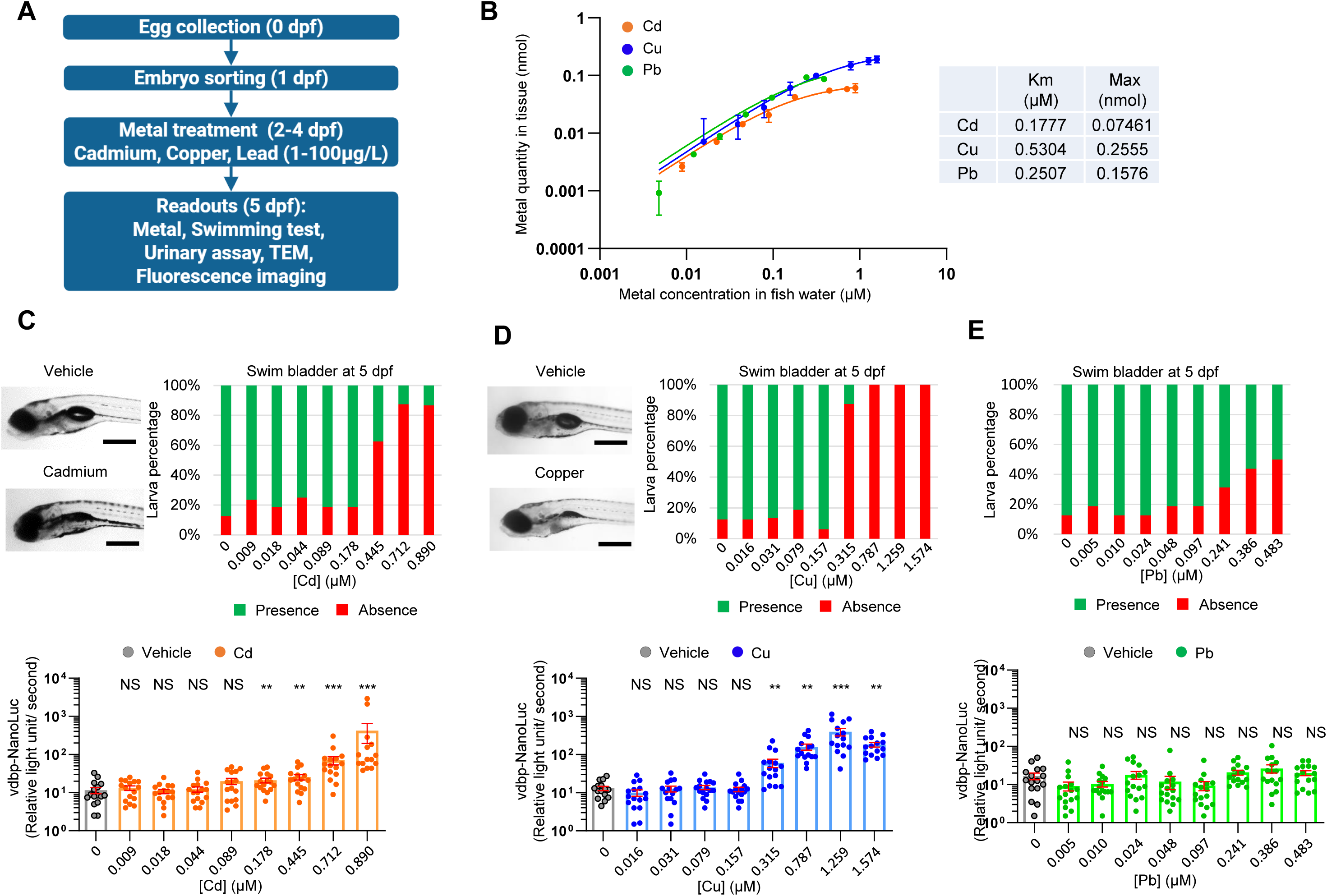
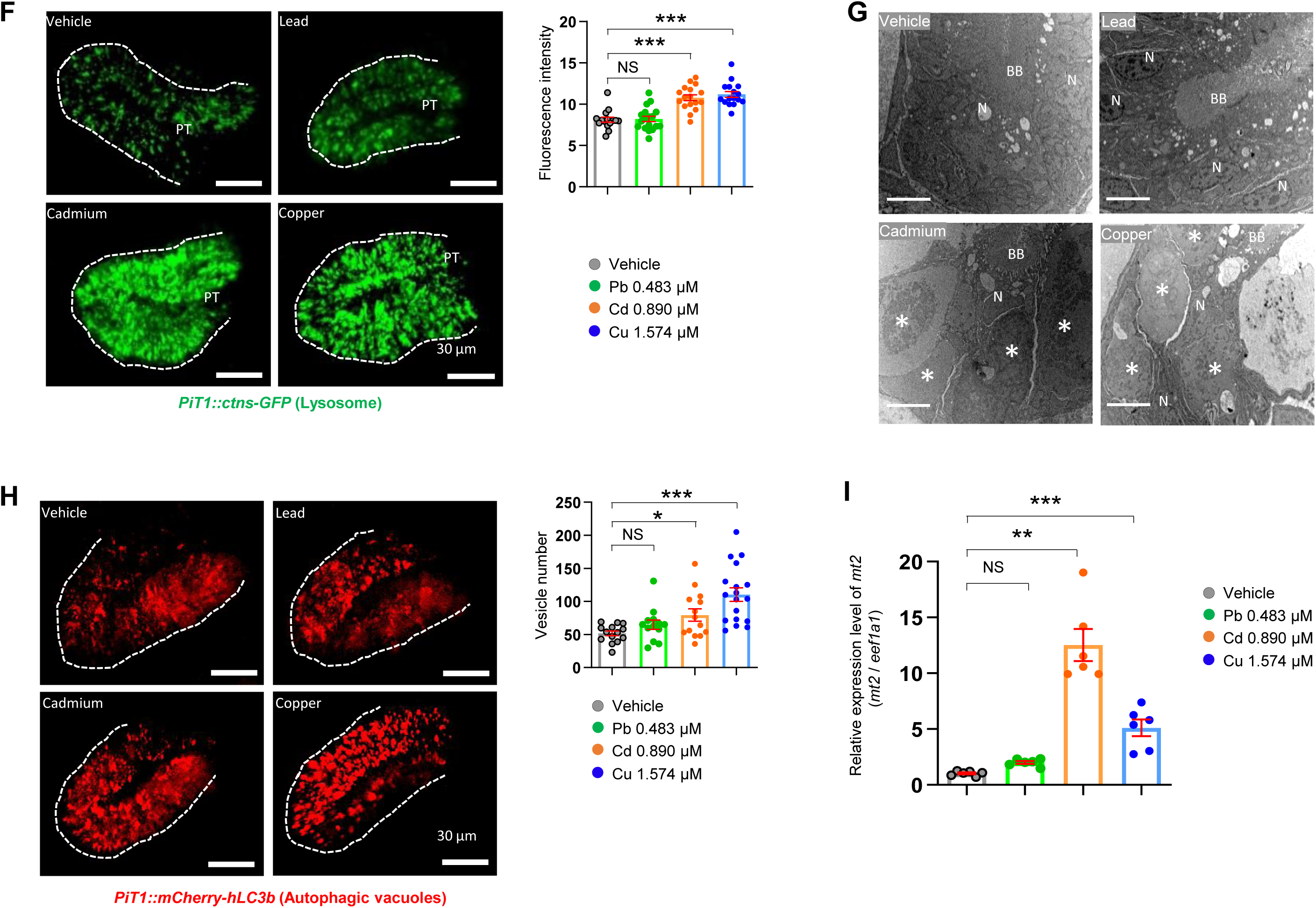
vdbp-NanoLuc reporter as biosensor for detecting metal nephrotoxicity. (A) Experimental protocol of metal toxicity assessment using *transgenic ½vdbp-NanoLuc* zebrafish larvae. (B) The quantity of Cd, Cu and Pb in larval lysate were analyzed by using Inductively coupled plasma mass spectrometry /ICP-MS after three days of incubation with metals. The concentration unit of metal was converted from µg/L to mM in order to be able to compare the concentration between different metals, as the molecular mass is different for the three metals. The Michaelis–Menten constant (K_M_) and maximal quantity in larvae (Max) are calculated from the measurement. The K_M_ of Pb is very similar to those of Cd, which are much lower than those of Cu, showing the lower affinity for uptake of Cu by larvae. (C) Semi-quantitative scoring of swim bladder defects and urinary analysis for 5-day-old larvae treated since 2 dpf with Cd at concentration from 0.009 to 0.890 µM. The swim bladder inflation was impaired by Cd treatment from the concentration of 0.445 µM. Urine samples were analyzed by luminometry. Excessive urinary loss of vdbp-NanoLuc was observed from the concentration of 0.178 µM. *n* = 16 (D) Semi-quantitative scoring of inflation defects of swim bladder and urinary analysis for 5-day-old larvae treated with Cu at concentration from 0.016 to 1.574 µM. Both defective inflation of swim bladder and excessive urinary loss of vdbp-NanoLuc were observed in larvae treated by Cu from the concentration of 0.315 µM. *n* = 16 (E) Semi-quantitative scoring of swim bladder inflation defects and urinary analysis for 5-day-old larvae treated with Pb at concentration from 0.005 to 0.483 µM. No LMW proteinuria was seen in any Pb-treated group. *n* = 16 (F) Microscopic examination of 5-day-old *Tg(PiT1::ctns-EGFP)* zebrafish larvae expressing lysosomal marker ctns-EGFP in PT of larvae treated with vehicle, Pb, Cd or Cu by using multiphoton fluorescence microscopy. Accumulation of ctns-EGFP-positive vesicles were accumulated in PT of larvae treated with 0.890 µM Cd or 1.574 µM Cu. PT: proximal tubule; *n* = 15 (vehicle), *n* = 18 (Pb), *n* = 17 (Cd), *n* = 16 (Cu); Bar = 30 µm. (G) The ultrastructure of PT in 5-day-old larvae treated by vehicle, Pb, Cd, and Cu was analyzed by transmission electron microscopy. The large electron-dense vesicles corresponding to the lysosome were accumulated in PT epithelial cells of larvae treated with 0.890 µM Cd or 1.574 µM Cu. Asterix: large electron dense vesicles; BB: brush border N: nucleus. Bar = 5 µm. (H) Microscopic examination of 5-day-old *Tg(PiT1::mCherry-hLC3b)* zebrafish larvae expressing autophagy marker mCherry-hLC3b in PT of larvae treated with vehicle, Pb, Cd or Cu by using multiphoton fluorescence microscopy. mCherry-hLC3b-positive autophagic vacuoles were accumulated in PT of larvae treated with 0.890 µM Cd or 1.574 µM Cu. PT: proximal tubule; *n* = 14 (vehicle), *n* = 13 (Pb), *n* = 14 (Cd), *n* = 18 (Cu); Bar = 30 µm. (I) Analysis of metallothionein *mt2* mRNA expression levels in metal-treated larvae by quantitative PCR. Total mRNA were extracted from larvae treated by vehicle, 0.483 µM Pb, 0.890 µM Cd or 1.574 µM Cu. After reverse transcription, the cDNA was analyzed by quantitative PCR to assess the *mt2* mRNA expression level. *n* = 6. Plotted data represent mean ± SEM. Nonparametric Mann-Whitney test, *P < 0.05, **P < 0.01, ***P < 0.001; NS, non-significant

The uptake of metal contaminants was monitored by ICP-MS analysis of larval lysate, yielding Michaelis–Menten kinetics (Fig. 4B). The uptake kinetic for the three cations were relatively identical, with *K*_m_ ranging from 0.2 (for Cd and Pb) to 0.5 **μ**M for Cu, however the saturation increase in the order Cd < Pb < Cu. Exposure to 0.178 µM Cd led to significantly higher urinary vdbp-NanoLuc levels, whereas exposure to 0.445 to 0.890 µM of Cd impaired swim bladder inflation in zebrafish larvae (Fig. 4C). Similarly, treatment of larvae with 0.315 to 1.574 µM of Cu led to increased urinary vdbp-NanoLuc and swim bladder defects (Fig. 4D). In contrast, Pb exposure led to no detectable increase in urinary vdbp-NanoLuc levels and much milder anomalies in swim bladder inflation (Fig. 4E). The total swimming distance was not affected by treatment with Cd and Cu (Fig. S3A,B) but significantly declined in groups exposed to 0.097-0.483 µM of Pb (Fig. S3C). The improved sensitivity of the vdbp-NanoLuc reporter system was verified for the Cu toxicity model, compared to the vdbp-mCherry system (Fig. S4).

Previous work has shown that Cd exposure impairs lysosome function *in vitro* (Fotakis et al., 2005), and that lysosomes play an important role in Cu homeostasis (Polishchuk and Polishchuk, 2016). To further investigate the mechanism by which metal contaminants yield differential PT toxicity, we assessed lysosome morphology and function using the PT-specific lysosome reporter *Tg(PiT1::ctns-EGFP*) (Chen et al., 2020) and autophagy reporter *Tg(PiT1::mCherry-hLC3b)* lines (Berquez et al., 2023; Nieri D, 2025) Treatment with Cd and Cu - but not with Pb - induced the accumulation of ctns-EGFP-positive lysosomes in PT cells (Fluorescence intensity, Cd:10.8 ± 0.4 vs. Cu: 11.2 ± 0.4 vs. Pb: 8.2± 0.3 vs. control 8.1±0.3; Fig. 4F), also reflected by the accumulation of enlarged electron-dense vesicles (Fig. 4G). These changes induced by Cd and Cu were paralleled by the accumulation of mCherry-hLC3b-positive autophagic vacuoles in PT cells, as compared to the Pb or vehicle-treated larvae, substantiating a defective lysosomal processing (Fig. 4H). The expression of metallothionein (mt2), a 7 kDa cysteine-rich metal-binding protein, can be induced by Cd in zebrafish larvae (Chen et al., 2004; Sabolic et al., 2010; Wu et al., 2008). We confirmed that exposure to these-cations induced a 12-fold (Cd) and 5-fold (Cu) increase in the expression level of *mt2* mRNA in the *½vdbp-NanoLuc* larvae, while no significant change was observed in the Pb-treated group, as compared to vehicle-treated controls (Fig. 4I).

### Inter- and intra-assay variability of the vdbp luminometric assay

The consistency and precision of bioluminescence measurement within and between different runs are essential for the capacity of the vdbp-NanoLuc system to screen compounds. By analyzing the urine samples collected from *lrp2a KO* or gentamicin-treated larvae, the luminometric assay showed low intra-assay variability, with a coefficient of variation (CV) of 3.4% (*lrp2a KO*) and 3.0% (gentamicin) (Fig. S5A-C). The inter-assay CV was 9.8% (*lrp2a KO*) and 8.2% (gentamicin) (Fig. S5A-C). The high intra-assay repeatability and inter-assay reproducibility confirmed the suitability of the vdbp-NanoLuc reporter system for drug screening for the genetic and toxic models used.

### Use of the vdbp-NanoLuc reporter system for nephron-protective drug screening

We next tested the value of the vdbp-NanoLuc system combined with an automated assessment of larval swimming behavior using the Zebrabox tracking system to test for potential compounds able to counteract cisplatin nephrotoxicity (Fig. S6; Fig. 5A). As oxidative stress is involved in the pathogenesis of cisplatin-induced nephrotoxicity (Soni et al., 2018), we selected and tested a library of 30 antioxidant compounds including FDA-approved drug, natural molecules, or molecules classified as Generally Recognized as Safe (GRAS) for their nephroprotective effect against cisplatin-nephrotoxicity (Table S1). To determine non-lethal concentrations, the *½vdbp-NanoLuc* larvae were treated from 2 dpf to 5 dpf at a maximal concentration of 1000 µM for water-soluble compounds, and 10-100 µM for water-insoluble compounds. Non-lethal concentrations were then used to screen for the rescue of cisplatin toxicity.

**Fig. 5.**
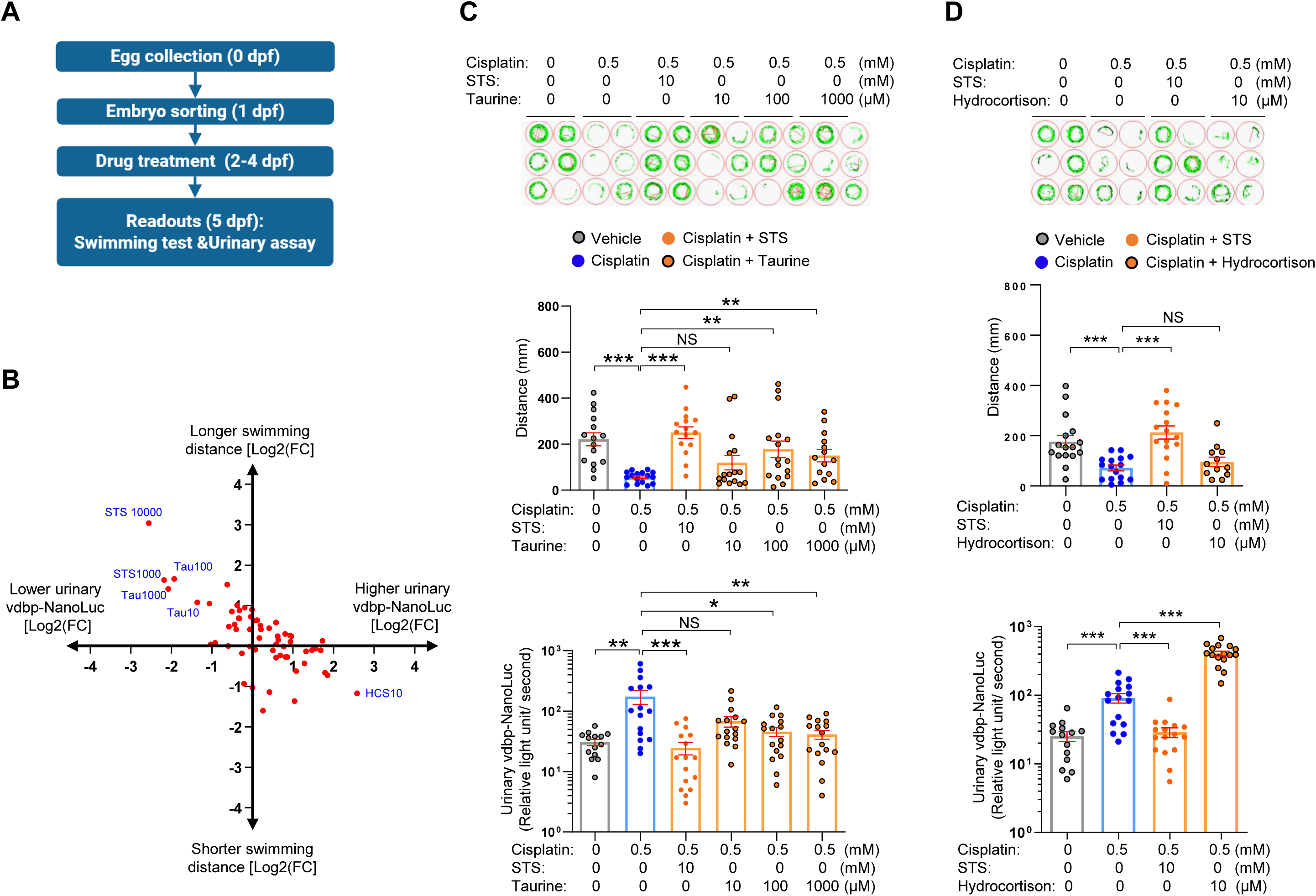
Use of the vdbp-NanoLuc reporter system for nephroprotective drug screening. (A) Experimental protocol of drug screening using transgenic *½vdbp-NanoLuc* larvae to identify antioxidants able to rescue the cisplatin nephrotoxicity (B) Graphical overview of screening data showing the log2-fold change in both urinary vdbp-NanoLuc reflecting the LMW proteinuria (X-axis) and swimming distance (Y-axis). Sodium thiosulfate (STS) at 10 mM (STS 10000) is able to completely prevent the cisplatin toxicity with strong improvement of swimming distance and reduction in urinary vdbp-NanoLuc luciferase activity. Urinary vdbp-NanoLuc luciferase activity was partially rescued by co-treatment with 100 µM and 1000 µM taurine (Tau100 and Tau1000), but increased by co-incubation with 10 µM HCS. Blue: drug name and concentration (µM). (C) Behavioral analysis and urinary analysis for 5-day-old zebrafish larvae incubated with vehicle, cisplatin alone, or co-incubated with cisplatin and 10 mM STS or taurine at different concentrations for 24 hours. *n* = 16. Overnight urine were collected and analyzed by luminometry; *n* = 16. (D) Behavioral analysis and urinary analysis for larvae incubated with vehicle, cisplatin alone, or co-incubated with cisplatin and 10 mM STS or 10 µM hydrocortisone (HCS) for 24 hours. Swimming capacity was not further damaged by HCS co-treatment, while urinary vdbp-NanoLuc luciferase activity was highly increased by HCS co-treatment. *n* = 16. Plotted data represent mean ± SEM. Nonparametric Mann-Whitney test, *P < 0.05, **P < 0.01, ***P < 0.001; NS, non-significant;

Using the combined phenotype analysis, STS at 10 mM (STS 10000) was able to completely rescue the LMW proteinuria and the swimming deficits caused by cisplatin alone, followed by taurine at both 100µM (Tau100) and 1000µM (Tau1000) (Fig. 5B). Conversely, treatment with hydrocortisone at 10 µM (HCS10) increased the urinary level of vdbp-NanoLuc (Fig. 5B). These protective and deleterious effects were confirmed by co-treatment of the *½vdbp-NanoLuc* larvae with STS, taurine or hydrocortisone. Both STS (10 mM) and taurine (100 μM and 1 mM) significantly rescued the swimming distance and the urinary vdbp-NanoLuc levels (Fig. 5C). The addition of hydrocortisone to cisplatin significantly increased the urinary vdbp-NanoLuc content, as compared to cisplatin alone (407±34 vs. 91.3±14 RLU/sec, respectively, p<0.01), which suggests a synergetic effect of hydrocortisone and cisplatin (Fig. 5D). Altogether, these data demonstrate the value of the *½vdbp-NanoLuc* larvae for drug screens by combining the bioluminescent urinary assay and swimming behavior analyses.

## DISCUSSION

Here, we establish a novel bioluminescent LMW proteinuria reporter system based on a transgenic zebrafish synthesizing the 43 kDa vdbp-NanoLuc protein in the liver. Our proof-of-concept studies indicate that the vdbp-NanoLuc reporter system can detect the LMW proteinuria (abnormal urinary excretion of vdbp-NanoLuc) associated with genetic deletion of endocytic receptor lrp2a/megalin or lysosomal cystine transporter cystinosin, or by exposure to nephrotoxic drugs (gentamicin and cisplatin) and metals (Cd and Cu). The reporter line allows to evidence the effect of small compounds on rescuing LMW proteinuria related to both genetic disorders and nephrotoxic drug exposure and to provide a screening protocol to identify nephro-protective compounds.

The zebrafish has emerged as a prominent model for studying human kidney disease and chemical nephrotoxicity due to its highly conserved nephron segment patterning (Outtandy et al., 2018; Poureetezadi and Wingert, 2016). The presence of LMW proteins in the urine is a consistent hallmark of PT dysfunction caused by either congenital or acquired disorders. A red fluorescence protein-based vdbp-mCherry reporter system was previously established and validated for monitoring LMW proteinuria and lysosomal processing in PT cells in zebrafish (Chen et al., 2020). As compared to ELISA assay of vdbp-mCherry, the quantification of vdbp-NanoLuc is more time- and cost-effective, with a 20-fold reduction in cost and 6-fold reduction in processing time for a 96-well plate (Table S2). The advantages of vdbp-NanoLuc quantification by luminometry also included higher sensitivity than the vdbp-mCherry by ELISA assay, and a low intra-assay and inter-assay variability.

A previous NanoLuc-based proteinuria reporter line (NanoLuc-D3) has been developed, based on the D3 domain of Receptor-Associated Protein (RAP), an endoplasmic reticulum-resident molecular chaperone that binds to megalin – and other members of the LDL receptor family (Naylor et al., 2022; Strickland et al., 2002). As NanoLuc-D3 is released from the liver into the circulatory system, the NanoLuc-D3 may interact with other LDL receptors and potentially affect their function. Moreover, as a chaperone fragment, the binding of D3 peptide to megalin might be mechanistically different from the binding of a bona fide, apical endocytic ligand to megalin. The bioluminescent vdbp-NanoLuc reporter system offers a more physiologically relevant model as endogenous VDBP produced by the liver serves as a significant marker of LMW proteinuria in both humans and rodents. Furthermore, 3 larvae per well has to be used when performing experiments with the NanoLuc-D3 system (Naylor et al., 2022), while a single larva per well is sufficient for all studies using*½vdbp-NanoLuc* zebrafish - highly simplifying larva handling for experimentation and compound screening.

Although cisplatin is an effective chemotherapy, its clinical use is limited by severe side effects including ototoxicity and nephrotoxicity. Acute kidney injury (AKI) is occurring in 20–30% of patients (Miller et al., 2010). Cisplatin nephrotoxicity has also been evidenced in zebrafish, including morphological alterations of PT cells and LMW proteinuria (Chen et al., 2020; Hentschel et al., 2005). Swim bladder defects were also observed when larvae were exposed to cisplatin. Sodium thiosulfate (STS), an antioxidant agent, is the sole FDA-approved medication used to treat cisplatin-induced ototoxicity. In rodent models, STS can alleviate cisplatin-induced toxicity, protecting treated animals from hearing impairment while reducing the levels of both blood creatinine and blood urea nitrogen (Dickey et al., 2005). Here, we provide evidence that STS treatment led to complete restoration of both swim bladder inflation and rescue of LMW proteinuria in zebrafish, providing insights into cisplatin-induced nephrotoxicity and potential therapeutic targets.

Water contamination by metals from the industrial and agricultural sector is a critical environmental issue (Lentini et al., 2017; Rawee et al., 2023). The kidney is a prime target of metal induced toxicity as the presence of transporters and endocytic receptors facilitates the uptake of metals into PT cells in particular (Rawee et al., 2023). Exposure of *½vdbp-NanoLuc* zebrafish larvae to environmentally relevant concentrations (from 1 to 100 µg/L) of Cd, Cu or Pb yielded specific toxicity patterns. Exposure of larvae to Cd and Cu dramatically impaired the inflation of the swim bladder and induced LMW proteinuria, contrasting with the very mild effects of Pb exposure on swim bladder inflation and the lack of PT dysfunction. The LMW proteinuria related to Cd and Cu exposure is associated with a marked accumulation of lysosomes in PT cells, as evidenced by a lysosome reporter line and TEM examination, and impairment of autophagy.

These changes were paralleled by a 12-fold (Cd) and 5-fold (Cu) increase in the expression level of *mt2* mRNA in the *½vdbp-NanoLuc* larvae. Previous work suggests that Cd-binding metallothionein (CdMT) is released from intoxicated hepatocytes, filtered and then endocytosed in the PT cells and degraded in lysosomes (Nordberg and Nordberg, 2022; Sabolic et al., 2010). The metal/metallothionein complexes was shown to associate with megalin, which mediate the uptake of these complexes by PT cells (Klassen et al., 2004; Wolff et al., 2006), thus the upregulated metallothionein *mt2* might drive the Cd- and Cu-induced kidney damage observed herein.

In contrast with Cd and Cu, Pb exposure did not induce PT dysfunction and LMW proteinuria; however it impacted on both larval development and swim behavior. Similarly, acute Pb exposure of freshwater rainbow trout did not induce excessive urinay protein excretion, but affected ionoregulation in the kidney (Patel et al., 2006). The mechanism by which Pb exposure causes toxicity in zebrafish larvae warrants further investigation. Since mild proteinuria has been detected in Wistar rats administered drinking water containing 1000 µg/L lead acetate for 8 weeks, resulting in a plasma Pb concentration of 355 µg/L (Missoun, 2010), species factors and/or differences in the type of Pb solution administered, dosage or duration could be involved. The effect of Pb exposure to reduced swimming distance could be a sign of neuronal impairment in the larvae and may be responsible for the mild defects observed in swim bladder inflation. Pb exposure induced neural alteration and neurotoxic symptoms in zebrafish larvae (Dou and Zhang, 2011; Roy et al., 2014). Of note, low-dose Pb exposure impairs early neuro-development, inducing lowered IQ, cognitive decline in infants (Lee and Freeman, 2014).

As a proof-of-concept, we performed a multiplex functional assay to screen for antioxidant compounds against cisplatin nephrotoxicity by combining the analysis of urinary vdbp-NanoLuc and larval swimming behavior. Our drug screens revealed the protective effect of taurine against cisplatin toxicity in zebrafish larvae: both urinary vdbp-NanoLuc and swimming activity were significantly rescued with 100µM and 1000µM taurine. This agrees with previous work in rodent models that found treatment with 150 – 250 mg/kg taurine prevents cisplatin cardiotoxicity (Chowdhury et al., 2016), brain injury (Owoeye et al., 2018) and gonadotoxicity (Azab et al., 2020). These results further support the use of zebrafish for *in vivo* small molecule screens.

Collectively, these studies demonstrate the reliability and value of the ½vdbp-NanoLuc biosensor in fish models of congenital and acquired dysfunction of the proximal tubule, including environmental contaminants such as metals. This novel system provides mechanistic insights into PT dysfunction and, when combined with a simple swimming test, a platform for drug screening to alleviate kidney tubular toxicity.

## MATERIALS AND METHODS

### Zebrafish maintenance

Zebrafish (*Danio rerio*) were kept at day/night cycle of 14/10 hours at 28°C in Fish Facility of University of Zurich (Zurich, Switzerland). Zebrafish eggs were obtained through natural spawning and raised in zebrafish E3 embryo medium containing 0.01% methylene blue. Larvae were fed with ZebraFeed 100-200 (Sparos, Portugal) beginning at 5 dpf. For experiments with *lrp2a* and *ctns* lines, knockout larvae were generated by crossing heterozygous lines. The experiments performed on animals were approved by the local legal authority (Veterinary Office, Canton of Zurich, Switzerland).

### Generation of transgenic Tg(lfabp::½vdbp-NanoLuc)

Multisite Gateway cloning technology was used to generate the expression vector *lfabp::½vdbp-NanoLuc/cmlc2::EGFP* using plasmids from Tol2 kit v1.2. The mCherry sequence in *pED-½vdbp-mCherry* (Chen et al., 2020) was replaced by NanoLuc luciferase cDNA (Promega, N1001) using BamHI / AscI restriction sites to create a middle entry clone *pED-½vdbp-NanoLuc*. The vector construction of *lfabp::½vdbp-NanoLuc/cmlc2::EGFP* was generated by LR Reaction using destination vector *pDestTol2CG2* with *cmlc2:EGFP* transgenesis marker, lab-generated 5’-entry clone *p5E-lfabp*, middle entry clone *pED-½vdbp-NanoLuc*, and 3’-entry clone *p3E-polyA*, as well as the Gateway™ LR Clonase™ II Enzyme Mix (Invitrogen, 11791-020, USA). Plasmid DNA at the concentration of 25 ng/µL was co-injected with Tol2 transposase mRNA at 50 ng/µL into wildtype zebrafish embryos at the one-cell stage to generate zebrafish with mosaic expression. Stable transgenic lines were generated by outcrossing founder zebrafish to AB wildtype fish. Primers used to clone NanoLuc cDNA are: *NanoLuc*-Fw: 5’-ACTGTTGGTAAAGGATCCCCGGTCTTCACACTCGAAGATTTC −3’, *NanoLuc*-Rev: 5’-TGCTCGAAGCGGCGCGCCGCCCCGACTCTAGA −3’. Double transgenic zebrafish larvae were produced by crossing transgenic *½vdbp-mCherry* zebrafish with transgenic *½vdbp-NanoLuc* zebrafish. For the analysis of mRNA expression level, full length of *½vdbp-NanoLuc* cDNA was amplified using primers vdbp-NanoLuc-Fw : ATGAATGCATCTTTAATTTTAATTTATGCTTTAATAGT and vdbp-NanoLuc-Rev : TTACGCCAGAATGCGTTCGC.

### Drug treatment

For the rescue of LMW proteinuria by rapamycin, *ctns* zebrafish larvae were incubated in fish facility water containing 200 nM rapamycin (MedChemExpress, HY-10219) from 5 dpf to 14 dpf, with daily refreshment of treatment conditions. At 14 dpf, overnight urine samples were collected for biochemical analysis.

For gentamicin treatment, 2-day-old embryos were injected with 1nL of 6 µg/µL gentamicin (Sigma, G1264) /0.9% NaCl solution alone or co-injected with 6.3 mg/mL taurine (Sigma, T0625) into the common cardinal vein after anesthesia with 0.2 mg/mL Ethyl 3-aminobenzoate methanesulfonate salt (Sigma, E10521). Embryos were immediately returned to fresh E3 medium until urinary analysis at 4 dpf.

The toxicity of cisplatin was assessed after incubating 2-day-old embryos in E3 medium containing 0.5 mM or 1 mM cisplatin (Sigma, C2210000) for 24 hours. Cisplatin was dissolved in E3 medium at room temperature with constant stirring for 6 hours. The rescue of cisplatin toxicity was performed using antioxidant drug sodium thiosulfate (STS) at 3 mM or 10 mM (Sigma, 72049). Treatment of 10 mM STS did not affect larva viability and pronephros morphology in healthy larvae (Al-Dahmani et al., 2022). Our pilot study showed that 10mM STS was well tolerated by wild type larvae over 3 days of consecutive treatment (from 2 dpf to 5 dpf). Larvae were co-treated with cisplatin and STS, followed by incubation in fresh E3 medium containing STS. Larvae treated only with vehicle or cisplatin, followed by incubation in E3 medium were used as controls. The inflation of the swim bladder at 4 dpf and mortality rate at 5 dpf were assessed by observation of larvae under an Olympus stereo microscope.

### Proof-of-concept drug screening to rescue cisplatin toxicity

The protocol used for drug screening is composed of two parts: determination of non-lethal concentration and rescue of cisplatin toxicity. Knowing that the sodium thiosulfate (water-soluble compound) could completely rescue LMW proteinuria at 10 mM concentration, working concentrations were initially fixed at 1000 µM, 100 µM and 10 µM for water-soluble compounds, which were directly dissolved in E3 or cisplatin/E3 solution. As cisplatin is not compatible with DMSO, water-insoluble compounds were first dissolved in N,N-Dimethylformamide/DMF (Sigma, 227056) and were tested at the concentrations of 100 µM, 10 µM and 1 µM. All compounds were purchased from Sigma.

The non-lethal concentration of each chemical was determined by incubation of wild type larvae in E3 medium with the chemical from 2 dpf to 5 dpf. Twenty larvae were tested for each condition using a 6-well microplate.

The protocol used to rescue cisplatin toxicity was designed in a way that can easily be adapted for large-scale screens, including automated movement tracking. The drug treatment, urine collection /analysis and movement tracking were performed in 96-well microplates. Transgenic *½vdbp-NanoLuc* embryos were sorted at 1 dpf by EGFP fluorescence in heart and dispensed in 96-well microplate with 1 embryo/well using EggSorter (Bionomous, Switzerland). At 2 dpf, larvae were treated with 200 µL of 0.5 mM cisplatin alone or cisplatin supplemented with the antioxidant (sodium thiosulfate or other drug candidates). 24 hours after cisplatin exposure, 150 µL of treatment solution was removed and replaced with 150 µL of fresh E3 medium alone or antioxidant-containing E3 medium. The larvae were then incubated for 48 hours with daily refreshment of the treatment solution. 50 µL of urine-containing E3 medium was collected from each well at 5 dpf for urinary analysis by luminometry. The 96-well microplate containing the larvae and remaining 150 µL of treatment solution was then used to track fish movement over a 5-minute swim trial using the Zebrabox (ViewPoint, France) to evaluate the behavior and total swim distance for each larva. We used larvae that were co-treated with cisplatin/sodium thiosulfate as a positive control and larvae treated with cisplatin alone as a negative control for each 96-well microplate. Co-treatments with cisplatin/antioxidant that killed more larvae than those treated with cisplatin alone were excluded from further testing. Drug effects were evaluated by comparing the cisplatin/antioxidant-treated zebrafish with those treated with cisplatin alone for both urinary and behavioral analyses. The data obtained were graphically represented with a log2-fold change in both quantification of NanoLuc luciferase activity/LMW protienuria (relative light intensity, X-axis) or total swimming distance (Y-axis).

### Metal exposure and analysis by Inductively coupled plasma mass spectrometry (ICP-MS)

To conduct our exposure to dissolved trace metals, plastic wares were pre-washed, with acid and ultrapure baths. The stock solution used in this study were 1001mg/L Cu(NO3)2 in 2% HNO3, (Copper standard for ICP, Fluka), 0.1 M Cd(NO3)2 (Cadmium ion standard, Fluka), 0.1M Pb(NO3)2 (Lead ion standard, Fluka). Intermediate solution of 100 mg/L Cu(NO_3_)_2_, Cd(NO_3_)_2_ or Pb(NO_3_)_2_ were prepared in 1% nitric acid, and further diluted in E3 medium for a working solution of 100 µg/L.

Following pH adjustment using NaOH, the 100 µg/L metal solution was used to prepare a series of treatment solutions with concentrations ranging from 1 to 80 µg/L of metal prepared in E3 medium. For both treatments in 96-well plate (with 1 larva/well, urinary analysis) or in 12-well plate (with 10 larvae/well, for metal quantification in larval lysate), 200 µL of metal solution per larva was used.

Following metal exposure, 10 larvae for each group were collected and placed in pre-washed 1.5 mL tubes. After removal of excessive treatment solution, larval tissues were transferred into new 1.5 mL tube and stocked at −80°C until sample processing after snap freezing in liquid nitrogen.

Larvae were digested using 100 µL of 65 % nitric acid and left at room temperature overnight. Larval samples were diluted with ultrapure water to reach a final concentration of 1% HNO_3_ and a volume of 10 mL. The concentrations of Cd, Cu and Pb were measured by using an ICP-MS (7700x, Agilent technology) under collision mode with He gas to avoid polyatomic interferences. Data obtained were used to investigate Michaelis-Menten uptake kinetic parameters for each metal salts.

### Fluorescence microscopy

A light sheet fluorescence microscope (ZEISS Light sheet Z.1, Germany) was used for *in vivo* visualization of EGFP expression in the heart of *½vdbp-NanoLuc* zebrafish larva using the 20x/1.0 NA water immersion objective (W Plan APO, Zeiss).

The Leica SP8 MP DIVE FALCON multiphoton microscope was used for high-resolution imaging of ctns-EGFP (lysosome marker), and vdbp-mCherry (endocytic ligand) in proximal tubule cells using the HC IRAPO L 25x/1.0 W motCORR objective lens and excitation by 2-photon laser at 920 nm (EGFP) or 1040 nm (mCherry). Zebrafish larvae bred into the nacre background were used for experimentation and collected after tail removal (the last third of zebrafish body length) followed by fixation in 4% PFA for overnight at 4°C. After three washes with PBS containing 0.1% Tween, zebrafish samples were embedded in 1% low-melting-temperature agarose for 3D optical imaging. The images acquired with multiphoton microscopy was processed with Huygens (Scientific Volume Imaging) for deconvolution. Total vesicle number in each pronephric PT was automatically quantified using Imaris software (Bitplane) and total fluorescent intensity was quantified using ImageJ and presented as relative fluorescent intensity.

### Bioluminescence microscopy

Visualization of bioluminescent vdbp-NanoLuc protein in transgenic zebrafish larvae was carried out at the Photonic Bioimaging Center at the University of Geneva. Five-day-old *½vdbp-NanoLuc* larvae were immobilized using MS222 and incubated in E3 medium containing the NanoLuc luciferase substrate, furimazine (1µL furimazine /100 µL E3 medium) and observed using an LV200 microscope (Olympus, Germany).

### Transmission electron microscopy

*lrp2a* larvae and *ctns* larvae were harvested at 5 dpf and chemical treated larvae at 14 dpf for ultrastructural analysis. The tails of the larvae were removed before overnight fixation in 2.5 % glutaraldehyde and 1.6 % paraformaldehyde in 0.1M pH7.3 cacodylate buffer. Samples were then rinsed 3x in 0.1M cacodylate buffer, post-fixed in 1 % osmium tetroxide/ cacodylate buffer for 40 minutes and stained in 1 % aqueous uranyl acetate for 1 hour. Dehydrated through a graded series of ethanol solution, larvae were infiltrated in 50% Epon812/ Propylene Oxide overnight, followed by embedding in Epon812 at 60°C for 28 hours. 350nm semi-thin sections were prepared using a Leica EM FCS ultra-microtome (Leica Microsystem, Germany), stained using a Toluidine blue solution, and examined under a light microscope. 60 nm ultra-thin sections were collected onto formvar-coated copper grids, stained with lead phosphate aqueous solution, and visualized using an electron microscope (FEI Tecnai G2 Spirit) at 120 kV. ImageJ software was used to analyze the TEM images.

### Tissue isolation and urine collection

Liver and kidney tissue were extracted from transgenic adult zebrafish in Nano-Glo® Luciferase Assay Buffer (Promega, N1120) and sonicated with a Digital Sonifier (Branson, 102C). Individual, overnight urine samples were collected from 5 dpf or 6 dpf larvae cultured in a 96-well plate with 100 μL or 200 μL E3 medium/larvae/well, respectively. At 14 dpf, 500 μL fish facility water was used to culture individual larvae for 16 h at 28.5°C to collect the overnight urine for both *½vdbp-mCherry* and *½vdbp-NanoLuc*. 50 µL fish pool water was then collected for biochemical analysis via ELISA or luminometry. Fish pool water samples for −20°C until the ELISA analysis could be carried out. Freshly collected fish pool water or tissue lysate was assessed immediately for NanoLuc luciferase activity.

### mCherry ELISA

Urinary ½vdbp-mCherry was quantified by one step assay using the mCherry ELISA kit (Abcam, ab221829). 50 µL of fish pool water was mixed with 50 µL of solution containing the capture antibody and detector antibody using a 96-well microplate. Samples in each well were then incubated at 37°C for 1 hour, rinsed 3x with Wash buffer PT, and incubated with 100 μL of TMB Substrate at room temperature for 10 minutes. The reaction was stopped by adding 100 μL of stop solution into each well and the absorbance of the reaction solution was read at 450 nm.

### Bioluminescent assay

#### Assay protocol

The NanoLuc luciferase activity was assessed in 96F non-treated, white microwell plates (Thermo Scientifc, 236108) for both luciferin furimazine (Promega, N1120) and Hikarazine™ Z108 (Synthelis Biotech, France) (Coutant et al., 2020). To prepare the Hikarazine Z108 solution, 1 mg of Hikarazine Z108 powder was dissolved in 0.2 mL DMSO, followed by the addition of 0.3 mL of acidic ethanol, which is prepared by adding 100 μL of 37 % hydrochloric acid in 12 mL of 100 % ethanol. The solution was incubated for 2 hours in a 50°C water bath then stored at −20°C until the time of assay. A custom-made NanoLuc assay buffer (NAB) was prepared for the urine assay using Hikarazine Z108. NAB buffer consisted of 100 mM MES, pH 6.0, 1 mM CDTA, 0.5% Tergitol NP-40, 0.05% Antifoam 204, 150 mM KCl, 1 mM DTT, and 35 mM thiourea (Hall et al., 2012; Xiong et al., 2022). 50 μL of fish pool water or tissue/larval lysate was assayed with 50 μL substrate/buffer mix for both furimazine and Hikarazine Z108. E3 medium/ fish facility water incubated alone (without larvae) was used as blank for normalization. The bioluminescence signal was measured using an Infinite M Plex Microplate reader (TECAN, Switzerland). Each microplate was read by the luminometer 3-times over the 10 minutes following the addition of substrate/buffer mix.

#### Optimization of substrate dosage for bioluminescent assay

In order to optimize our protocol of bioluminescent assay, we tested two different concentrations of substrate for both furimazine and Hikarazine Z108 using larval lysate containing a small amount of vdbp-NanoLuc protein. We conducted a serial dilution of larval lysate extracted from *½vdbp-NanoLuc* larvae in fish E3 medium using 1µL (dose suggested by Promega) or 0.5 µL of furimazine per assay. The light intensity obtained from the reactions using 0.5 µL of furimazine is comparable to the values assayed with 1 µL (Fig. S1B), demonstrating that 0.5 µL furimazine per assay is enough to obtain an optimal light intensity in our system. We also tested two concentrations of Hikarazine Z108: 0.31 µL/assay (with final concentration of 13 µM, as suggested by Coutant *et al*.), or 0.5 µL /assay (with final concentration of 21 µM). The assay with 0.5 µL Hikarazine Z108 displayed higher light intensity than those of 0.31 µL (Fig. S1C). The light intensity is directly proportional to the relative concentration of NanoLuc luciferase for both furimazine and Hikarazine Z108 (Fig. S1B,C). Next, we compared the light intensity given by the bioluminescent reaction using furimazine and Hikarazine Z108 with 0.5 µL /assay. Overall, the light signals obtained with Hikarazine Z108 were approximately three times greater than the values assayed with furimazine (Fig. S1D). However, the half-life time of light obtained with Hikarazine Z108 (around 60 min) is shorter than those in assay done with furimazine (around 90 min). Due to the difference in half-life time, we decided to use furimazine in the remainder of the analyses presented here, except for the screening experiment.

### Quantitative real-time PCR

Ten zebrafish larvae were homogenized in 1mL PureZOL™ RNA Isolation Reagent (Bio-Rad, 732-6880, USA) using T10 Basic Ultra-Turrax Disperser (IKA, Staufen, Germany). Total RNA was extracted and purified by Aurum Total RNA Fatty and Fibrous Tissue Kit (Bio-Rad, 732-6830, USA). Genomic DNA was removed by on-membrane DNAse I treatment during RNA purification. 1000 ng of RNA was converted into cDNA by reverse transcription with iScript cDNA Synthesis Kit (Bio-Rad), followed by determination of mRNA expression levels using relative transcript quantification by Quantitative PCR with iQ SYBR Green Supermix (Bio-Rad) and CFX96 Real-Time PCR Detection System (Bio-Rad). Quantitative PCR primers were designed on the website of Primer3. The sequences are zf-eef1a1a-Fwd : TTCTCCGAGTATCCTCCTCTG, zf-eef1a1a-Rev : CTTCTCCACTCCTTTAATCACTCC, zf-mt2-Fwd: CCTCCAGCATCAACTCATTCAC, zf-mt2-Rev: CAACTCTTCTTGCAGGTAGTACAC. PCR temperature cycling conditions were 95°C for 3 min followed by 40 cycles of 15 s at 95°C and 30 s at 60°C. The relative changes in mRNA level of target genes over housekeeping gene *eef1a1a* were calculated using the Delta-Delta Ct Method.

### Western blot

Proteins were extracted from liver isolated from adult zebrafish or larval tissue using RIPA buffer (Sigma, R0278) supplemented with 20% glycerol and after sonication with Digital Sonifier (Branson, 102C). Protein concentration was determined using Bradford protein assay (BioRad, 5000002). Proteins were separated by SDS-PAGE under reducing conditions and transferred onto PVDF membrane. Non-specific binding sites were blocked by incubation in 5% non-fat milk (Bio-Rad, 1706404) diluted in PBS. The membrane was incubated overnight at 4°C with primary antibody, then peroxidase-labeled secondary antibody (Darko, Denmark). The peroxidase activity was assayed with Immobilon Western Chemiluminescent HRP Substrate (Millipore, WBKLS0500) and visualized with ChemiDoc™ Touch Imaging System (Bio-Rad). The following antibodies were used in this study: anti-NanoLuc luciferase (Promega, N7000) or anti-gapdh (Cell signaling, 14C10), anti-GFP (Thermo Fisher, A-11122).

### Statistical analysis

Statistical analyses were performed using Prism 5 software (GraphPad Software, USA). A D’Agostino-Pearson normality test was applied on all data sets. Either unpaired two tailed t-test or Mann-Whitney non-parametric tests were used to compare treatment groups. Zebrafish embryos were randomly selected for chemical-treatment group or control group. Statistical significance was set at a p<0.05.

## Acknowledgements

We acknowledge Dr. Christoph Ruediger Bauer, Dr. Andres Käch and Zsuzsa Radvanyi for providing technical assistance. Dr Yves L. Janin is acknowledged for kindly providing Hikarazine Z108 samples. We also thank zebrafish caretaker team of zebrafish facility at the University of Zurich. Fluorescence imaging and analysis with transmission electron microscopy was performed with equipment maintained by the Center for Microscopy and Image Analysis at the University of Zurich. Bioluminescence images were acquired at the Photonic Bioimaging Center of the University of Geneva.

## Author contributions

Conceptualization: Z.C., A.L., O.D.,V.I.S.; Methodology: Z.C., I.W.; Resources: O.D., S.C.F.N., V.I.S.; Formal analysis: Z.C., H.L., I.W.; Investigation: H.L., M.G., I.W., S.K.; Writing - original draft: Z.C., O.D.; Writing - review & editing: A.L., S.C.F.N., V.I.S.

