## Supplementary figures and tables for "A Novel Zebrafish Luminescent Biosensor for Kidney Tubulopathy, Metal Toxicity, and Drug Screening"

### <u>Supplementary material</u>

Supplementary Fig. S1 to S6

Supplementary Tables S1 and S2

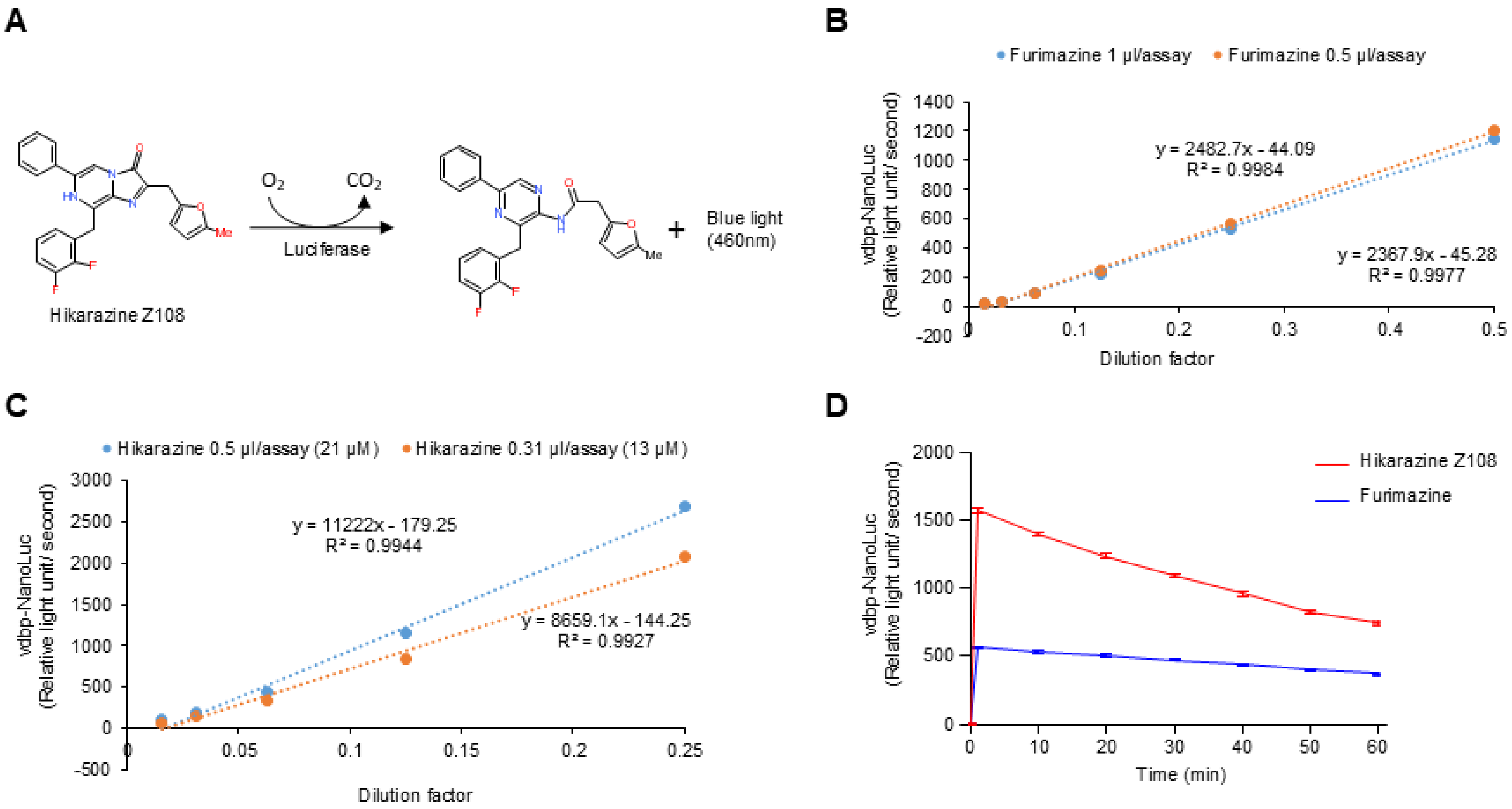


**Fig. S1: Luciferins and optimization of NanoLuc luciferase assay.**

(A) The oxidation of Hikarazine Z108 by NanoLuc luciferase in the presence of $O_2$ produces blue light of wavelength 460nm. (B) Bioluminescent VDBP signal was measured from NanoLuc luciferase assay performed with a serially diluted larval lysate containing ½vdbp-NanoLuc and its substrate furimazine. Two different doses of furimazine were employed for assay: 1 µl (dose suggested by Promega) or 0.5 µl furimazine in a final volume of 100 µl per assay; n = 4 replicates. (C) Bioluminescent VDBP signals were quantified from NanoLuc luciferase assay using a serially diluted larval lysate containing ½vdbp-NanoLuc and Hikarazine Z108 as substrate. Two different doses of Hikarazine Z108 were tested: 0.31 µl (dose suggested by Coutant *et al.*) or 0.5 µl Hikarazine Z108 in a final volume of 100 µl per assay; n = 4 replicates. (D) Dynamic changes of bioluminescent VDBP signal measured at different time points over a period of 60 minutes. Same amounts of larval lysate containing ½vdbp-NanoLuc were used in reactions for both furimazine and Hikarazine Z108; n = 8 replicates.

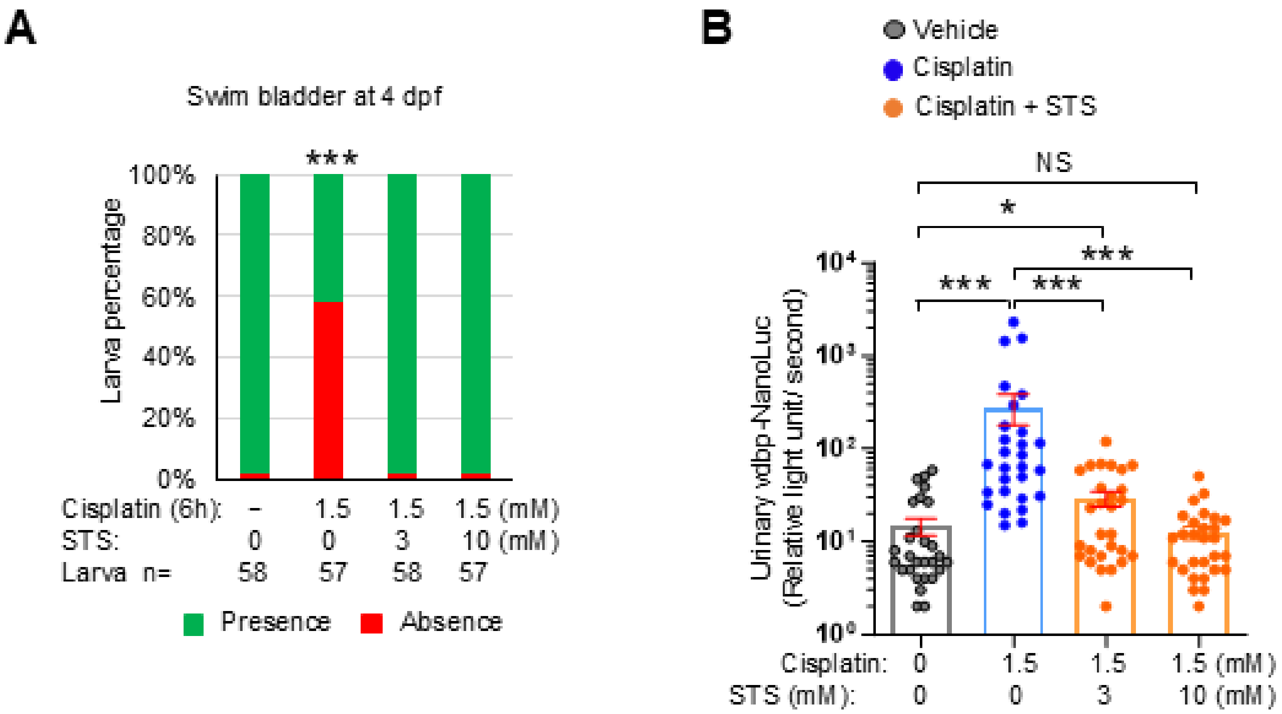


**Fig. S2. Rescue of cisplatin toxicity by co-treatment with sodium thiosulfate.**

(A) Morphological examination of 4-day-old zebrafish larvae treated with vehicle, 1.5mM cisplatin alone (6 hours) or co-treated with cisplatin/sodium thiosulfate (STS) at 2 dpf for 6 hours and semi-quantitative scoring of developmental defects of swim bladder ; Bar = 0.5 mm.

(B) Overnight urine were collected from 5-day-old transgenic *½vdbp-nanoLuc* zebrafish larvae treated with vehicle, 1.5mM cisplatin alone or co-treated with cisplatin /sodium thiosulfate (3 mM or 10 mM) at 2 dpf for 6 hours and urinary analysis by luminometry; *n* =28 (Vehicle), *n* = 28 (1.5 mM cisplatin alone) , *n* = 28 (1.5 mM cisplatin + 3 mM STS), *n* = 28 (1.5 mM cisplatin + 10 mM STS).

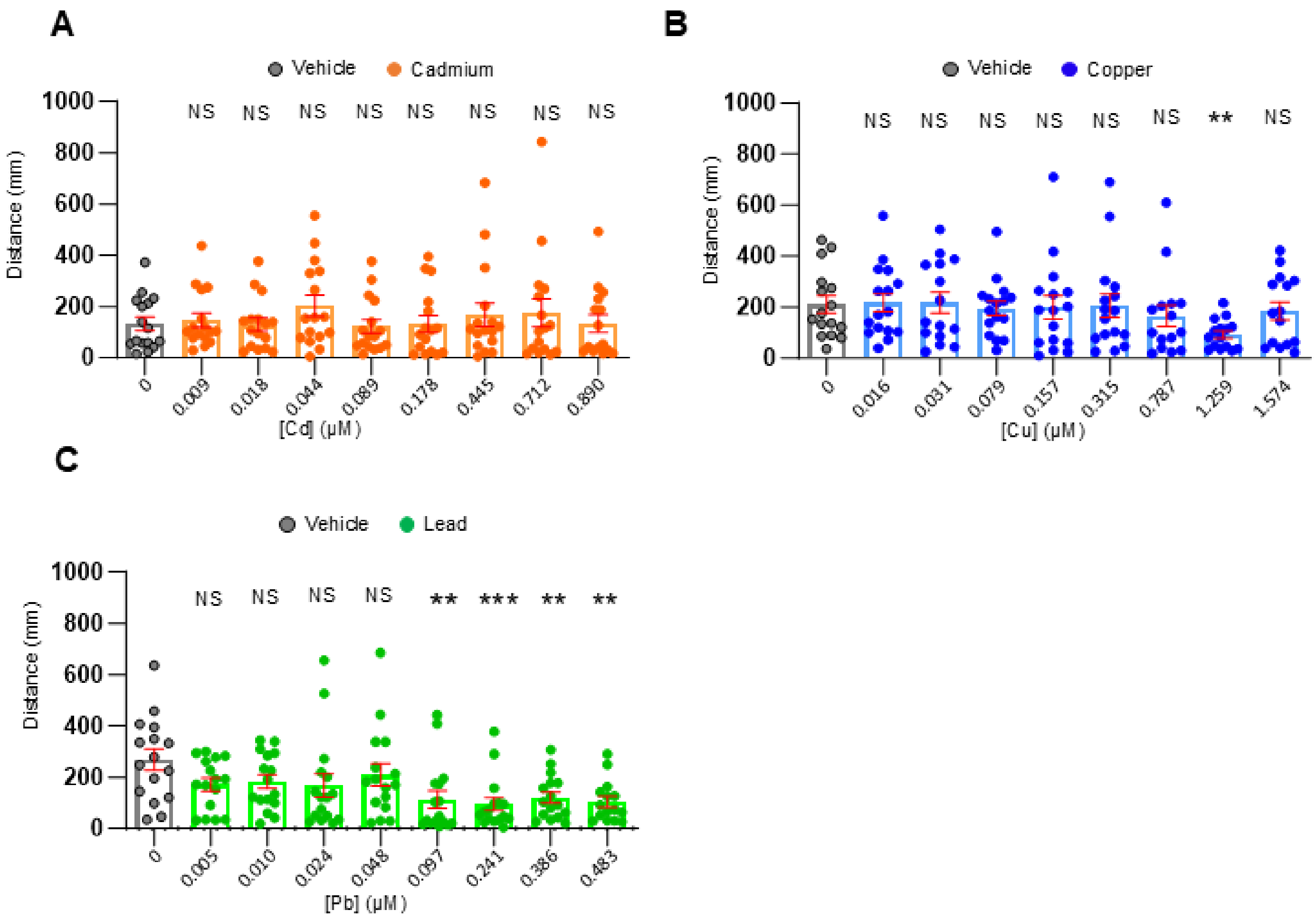


**Fig. S3. Assessment of swimming behavior in larvae treated with heavy-metals.**

(A) Behavioral analysis of 5 dpf zebrafish larvae incubated with vehicle or Cd at concentration from 0.009 to 0.890 µM. (B) Behavioral analysis of 5 dpf zebrafish larvae incubated with vehicle or Cu at concentration from 0.016 to 1.574 µM. Swimming distance impairment was only observed in larvae treated with 1.259 µM Cu. (C) Behavioral analysis of 5-day-old zebrafish larvae treated with vehicle or Pb at concentration from 0.005 to 0.483 µgM. Swimming distance declined significantly in groups treated with 20 - 100 µg/L of lead. *n* = 16.

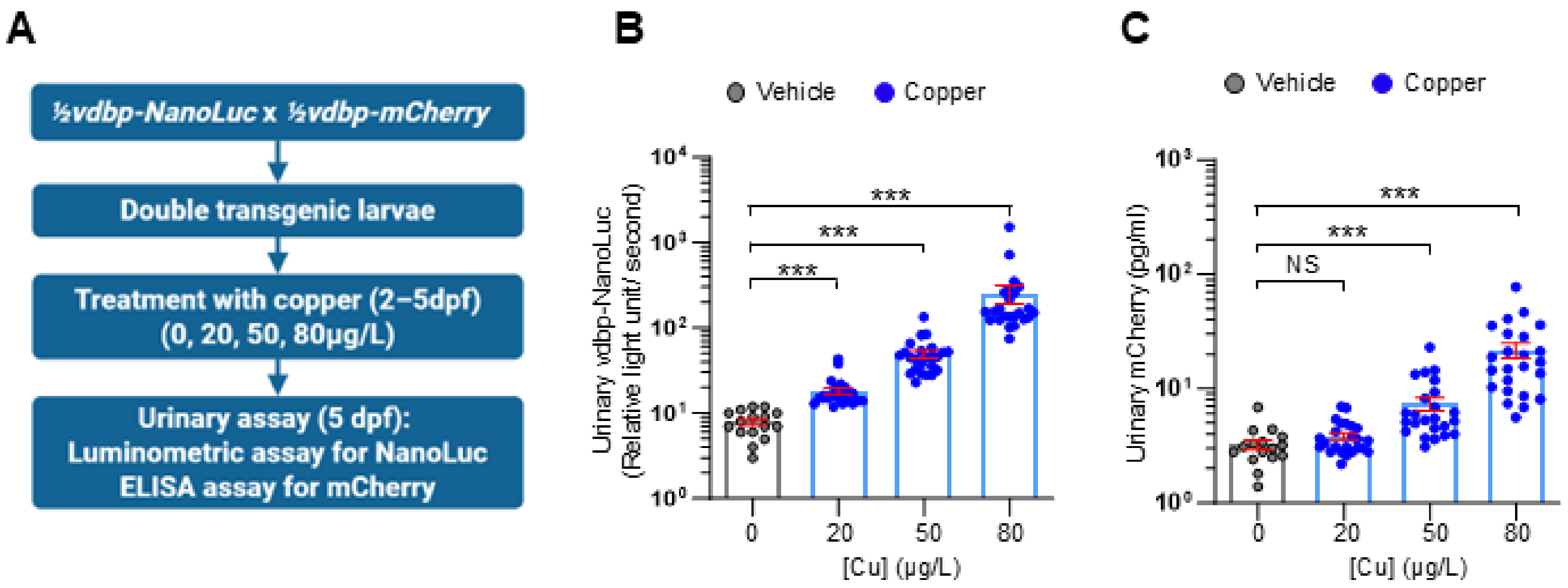


**Fig. S4: Comparative studies of LMW proteinuria evaluation using vdbp-NanoLuc and vdbp-mCherry in Cu toxicity models**

(A) Transgenic *½vdbp-NanoLuc* zebrafish was crossed with *½vdbp-mCherry* zebrafish to produce double transgenic larvae expressing both vdbp-NanoLuc and vdbp-mCherry tracers. Transgenic larvae were treated with Cu at the concentration of 0, 20, 50 and 80 µg/L. Urine samples were collected at 5 dpf and analyzed for both VDBP reporter proteins for the same urine sample. (B) The urinary NanoLuc luciferase activity was quantified by luminometric assay. The bioluminescent signals in all three Cu-treated groups are significantly higher than vehicle-treated control. n = 16 (vehicle), n = 24 (Copper-treated group). (C) The quantity of vdbp-mCherry in the same urine samples assessed in (B) was analyzed by mCherry ELISA assay. No difference was observed between the 20 µg/L Cu-treated group and vehicle-treated control. n = 16 (vehicle), n = 24 (Cu-treated group).

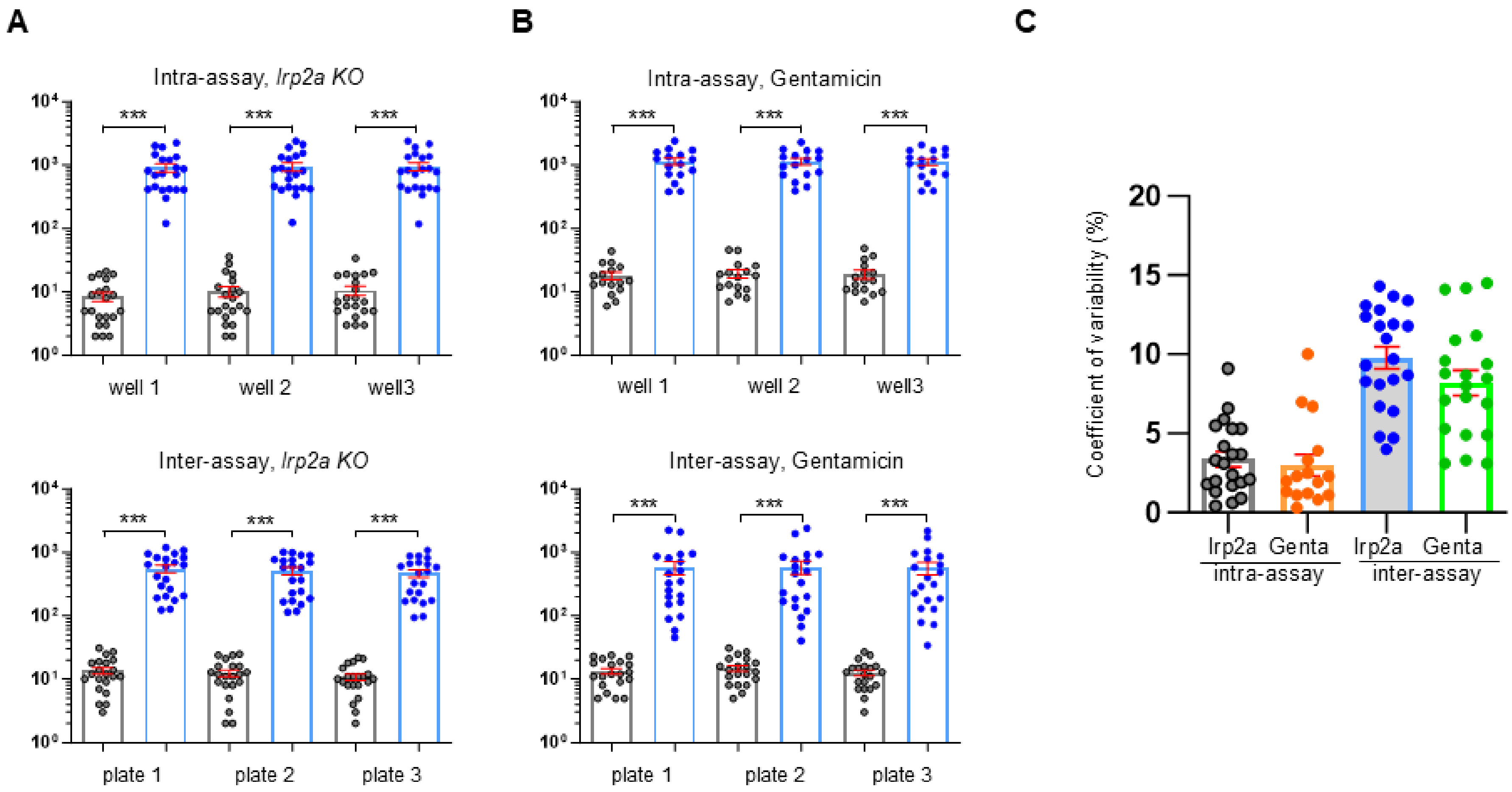


**Fig. S5: Inter- and intra-assay variability for quantification of vdbp-NanoLuc by luminometry**

(A) Urine samples collected from *lrp2a KO* larvae were analyzed by luminometry in triplicates (3 wells for each sample on a same plate, intra-assay variability) and in 3 different runs (1 well/plate and 3 plate, inter-assay variability). (B) Urine samples collected from gentamicin-treated larvae were analyzed by luminometry in triplicates (intra-assay variability) and in 3 different runs (inter-assay variability). (C) Calculation of intra- and inter-assay coefficient of variability (%) with data obtained from (A) and (B).

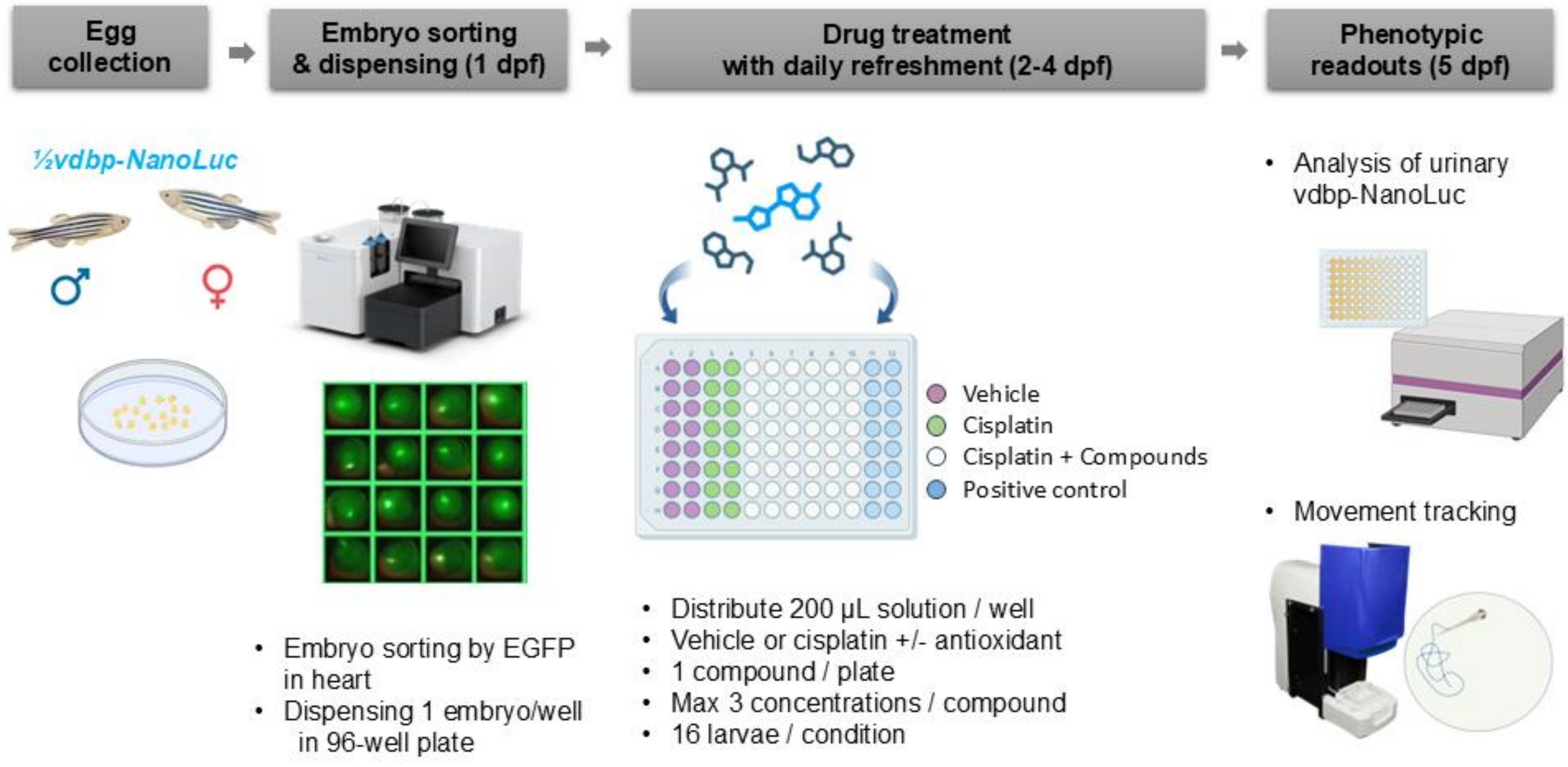


**Fig. S6. Experimental protocol of drug screening to antioxidants able to rescue the cisplatin-induced nephrotoxicity**

Bioluminescent *½vdbp-NanoLuc* larvae were treated with cisplatin and antioxidants from 2 dpf in 96-well plate and the treatment effect were assessed at 5 dpf. At 1 dpf: automated embryo sorting by EGFP fluorescence in heart and dispensing in 96-well plate; At 2-4 dpf: treatment of larvae with 200 µL cisplatin alone or mixture of cisplatin+ antioxidant solution, with daily refreshment of treatment; At 5 dpf: quantification of urinary vdbp-NanoLuc with 50 µl of urine / fish pool water and movement tracking of larvae kept in 150 µl of fish pool water in 96-well microplate.

| Name | Reference | | Vehicle | Solubility (mM) | Stock (mM) | 1000 µM | 100 µM | 10 µM | 1 µM |
|---|---|---|---|---|---|---|---|---|---|
| Aminothiazole (AMT) | 123129 | FDA approved | water | 1000 | 10 | OK | OK | OK | -- |
| Betaine (BTN) | B2629 | FDA approved | water | 13419 | 10 | OK | OK | OK | -- |
| Cysteamine (CYTM) | M6500 | FDA approved | water | 3521 | 10 | OK | OK | OK | -- |
| Gallic Acid (GLA) | G7384 | Natural | water | 70 | 10 | lethal | OK | OK | -- |
| N-Acetylcysteine (NAC) | A7250 | FDA approved | water | 613 | 10 | lethal | OK | OK | -- |
| Sodium thiosulfate (STS) | 72049 | FDA approved | water | 4427 | 10 | OK | OK | OK | -- |
| Taurine (Tau) | T8691 | GRAS | water | 500 | 10 | OK | OK | OK | -- |
| Dimethyl Fumarate (DMFM) | 242926 | FDA approved | DMF | 83.0 | 50 | -- | lethal | OK | OK |
| D-Pinitol (DPNT) | 441252 | Natural | DMF | 25.8 | 10 | -- | -- | OK | OK |
| Farnesol (FNS) | F203 | GRAS | DMF | 90.1 | 50 | -- | lethal | OK | OK |
| Ferulic acid (FRA) | PHR1791 | FDA approved | DMF | 103.1 | 100 | -- | lethal | OK | OK |
| Flavone (FVN) | F2003 | GRAS | DMF | 198 * | 100 | -- | lethal | OK | OK |
| Flavanone (FVNN) | 102032 | GRAS | DMF | 196 * | 100 | -- | lethal | OK | OK |
| Hesperidin (HSP) | H5254 | GRAS | DMF | 49.2 * | 50 | -- | OK | OK | OK |
| 4-Hydroxyphenylacetic acid (HPA) | H50004 | Natural | water | 329 | 10 | -- | OK | OK | OK |
| Hydrocinnamic acid (HCA) | 135232 | Natural | DMF | 200 * | 100 | -- | lethal | OK | OK |
| Hydrocortison (HCS) | H0888 | FDA approved | DMF | 82.9 | 100 | -- | OK | OK | OK |
| 3-Indolepropionic acid (IPA) | 220027 | Natural | DMF | 196 | 100 | -- | lethal | OK | OK |
| Melatonin (MLT) | M5250 | Dietary | DMF | 198 | 100 | -- | OK | OK | OK |
| Naringin (NRG) | 91842 | GRAS | DMF | 34.5 | 10 | -- | -- | OK | OK |
| Pyrogallol (PYG) | P0381 | GRAS | DMF | 238 | 100 | -- | OK | OK | OK |
| Resveratrol (RSV) | R5010 | Natural | DMF | 285 | 100 | -- | OK | OK | OK |
| Rosmarinic acid (RMA) | R4033 | Natural | DMF | 97 | 100 | -- | OK | OK | OK |
| Sesamol (SML) | S3003 | Natural | DMF | 217 | 100 | -- | OK | OK | OK |
| Silymarin (SLY) | S0292 | Natural | DMF | 41.5 | 10 | -- | -- | OK | OK |
| Sinapic acid (SNA) | D7927 | Natural | DMF | 44.6 | 10 | -- | -- | OK | OK |
| Syringic acid (SA) | S6881 | Natural | DMF | 80.8 | 50 | -- | OK | OK | OK |
| Syringaldehyde (SYA) | S7602 | Natural | DMF | 164.8 | 100 | -- | lethal | OK | OK |
| tert-Butylhydroquinone (TBHQ) | 112941 | Food additive | DMF | 198.8 * | 100 | -- | lethal | lethal | OK |
| Umbelliferon (UBL) | H24003 | Natural | DMF | 197.5 * | 100 | -- | OK | OK | OK |

**Table S1: List of antioxidants tested in this proof-of-concept screening and their non-lethal concentration determined in wild type zebrafish.**

Water-soluble compounds are tested with a maximal concentration of 1000 µM, while water-insoluble compounds are used at a maximal concentration of 100 µM. * Solubility in DMSO.

| | vdbp-NanoLuc | vdbp-mCherry |
|---|---|---|
| Protein size | 380 a.a. | 446 a.a. |
| Predicted MW | 43 kDa | 50 kDa |
| Detection method | Luminometry | ELISA |
| Cost for 96-well plate | **$ 35 (Furimazine)** | **$ 700** |
| Time for 96-well plate | **20 min** | **120 min** |
| Number of steps for assay | 3 | 8 |
| Assay procedure | Add 50 µl urine/Well | Add 50 µl urine/Well |
| | Add 50 µl assay buffer | Add 50 µl assay buffer |
| | Plate reading | Incubation 60 min |
| | -- | 3 x wash |
| | -- | Add 100 µl TMB solution |
| | -- | Incubation 10 min |
| | -- | Add 100 µl Stop solution |
| | -- | Plate reading |
| Inter-assay CV | < 10% | NA |
| Intra-assay CV | < 5% | NA |

**Table S2: Comparison of characteristics and detection procedure for vdbp-NanoLuc and vdbp-mCherry.** As compared to ELISA assay of vdbp-mCherry, the quantification of vdbp-NanoLuc allows a 20-fold reduction in cost and 6-fold reduction in processing time for a 96-well plate.
